# The fate of intracoelomic acoustic transmitters in Atlantic Salmon (*Salmo salar)* post-smolts and wider considerations for casual factors driving tag retention and mortality in fishes

**DOI:** 10.1101/2023.05.15.540815

**Authors:** M.J. Lawrence, B.M. Wilson, G.M. Reid, C. Hawthorn, G. English, M. Black, S. Leadbeater, C.W. McKindsey, M. Trudel

## Abstract

Acoustic telemetry is a widely used method in assaying behavioural dynamics in fishes. Telemetry tags are often surgically implanted in the coelom of the animal and are assumed to have minimal rates of post-release mortality and tag shedding. However, fish are capable expelling tags and mortalities do occur following release, with the mechanism(s) underlying these effects not well understood. The purpose of this research was to address causal factors underlying tagging mortality and tag expulsion in fishes. We conducted an empirical assessment of tag retention and post-surgical mortality rates in post-smolt Atlantic salmon (*Salmo salar*) fitted with a dummy acoustic tag over a 92 day monitoring period. This was complimented with a meta-analysis of factors affecting tag retention and post-surgical mortality rates in the wider literature. Post-smolt salmon exhibited low rates of mortality following tag implantation (≤ 5.1%) but had high rates of tag expulsion (54.8%) and impaired growth and a foreign body response was evident. The meta-analysis showed that mortality was generally low across all studies (12.4%) and was largely unaffected by model cofactors. Tag retention rates were high among the studies investigated here (86.7%) and had a weak negative relationship with tag:body mass ratios. Our results suggest that while mortality is often low among tagging studies, including this one, caution must be exercised in assessing stationary tags as they may represent an expelled tag rather than a mortality event. Our results also indicate that tag dimensions are not nearly important as the tag:body mass ratio.

## Introduction

Acoustic telemetry has become an increasingly popular method for assessing animal behaviour and spatial use patterns in contemporary aquatic biology [1,2]. At its core, acoustic telemetry involves the use of acoustic transmitters that are externally attached or internally implanted to an experimental animal, which relay position and can inform 0depth, acceleration, and/or environmental temperature of the focal organism to any receiver device in range [3]. The information provided by acoustic tags can be used to infer a wide range of life history characteristics and ecological interactions in aquatic organisms. These data can address specific questions pertaining to an animal’s daily and seasonal movement patterns [4–6], habitat preferences [7–9], and predation [10,11]. Overall, advances in acoustic telemetry have coincided with a renaissance in characterizing animal behaviour and ecology, furthering our foundational understanding of the biology of aquatic organisms.

In a fisheries context, acoustic telemetry has proven to be an important tool for stock management through development of an improved understanding of managed species’ biologies [1]. For example, telemetry has been used to assess rates of natural-, fisheries-, and predator-associated mortality in wild fishes [12–15], which can refine mortality model estimates in stock assessment profiles [1]. Furthermore, acoustic telemetry has provided great insight into the spatial use and migratory patterns of commercially-important species. In the case of Pacific salmonids, factors underlying migration timing and fine-scale migration patterns have been elucidated using telemetry to better understand the spatial ecology of these fishes [16–19]. Given the flexibility of acoustic telemetry, and the wealth of data it provides on numerous aspects of fish biology, it is well-suited for use in fisheries biology.

To characterize behaviours, fish are fitted with an acoustic transmitter, which is typically inserted into the celomic cavity through a small midventral incision that is sutured closed, followed by recovery and release of the tagged fish [20]. The underlying assumption of such tagging procedures is that the tag has a minimal effect on fish health and mortality and that the tag is retained in the body cavity until the end of the transmitter battery life. Indeed, there is support for this notion as, in a broad range of fish species, mortality and growth rates appear unaffected with tag retention levels being high [21–26]. Although prior work has established that post-tagging monitoring typically occurs over a rather acute duration (i.e. < 1 month [27]), longer-term impacts on mortality, growth, and tag retention are poorly understood. In addition, a comprehensive analysis of the effects of acoustic tags on animal recovery and tag fate (i.e. retention vs expulsion) has yet to be performed on fishes. Such considerations could be useful in making predictions of tag effects and fate in understudied systems.

To date, acoustic telemetry has been used to track the movements of a wide range of aquatic animal species including reptiles, invertebrates, and mammals [28]. Fishes, particularly the salmonids, remain the most studied taxa in this respect [28,29]. Despite widespread use of acoustic tags, there still exists a degree of uncertainty of how the tag may have adverse effects on the health and survival of the fish [30,31]. Indeed, there appears to be a dearth of information relating to the effects of acoustic transmitters on juvenile Atlantic salmon. From what little has been addressed, Atlantic salmon appear to have high levels of survivorship following tag implantation (upwards of 100% [21,32]), although studies using smaller sample sizes (n = 5) have noted that mortality can be upwards of 60% [33]. Furthermore, documentation of both abdominal tag expulsion and some basic characterisations of internal tag fate have also been made [21,32,,33]. Indeed, tag expulsion through the abdominal wall has been documented in salmon smolts [21,32,,33], with expulsion rates appearing to have a mass/size basis such that larger tags, relative to the fish, have lower retention rates [32,33]. Aside from these few works, virtually nothing is known about tagging-associated mortality and rates of tag retention in juvenile Atlantic salmon, hampering our ability to make interpretations surrounding the post-release fate of these fish in acoustic telemetry studies.

The purpose of this work is threefold: 1. Characterize post-surgical mortality in Atlantic salmon post-smolts over a timeframe reflective of a typical transmitter’s battery life (i.e. > 90 days), 2. assess the ultimate fate of an intracoelomically placed tag, with respect to tag retention rates and anatomical responses to the tag, and 3. conduct a review of the literature to address the impacts and fate of acoustic tags in a broad range of fish species to help develop a predictive framework associated with tagging effects. This exploratory work was conducted using two sizes of acoustic transmitters, the Innovasea V7 and V8, which are often used in Atlantic salmon telemetry studies [14,34,,35]. Fish were fitted with an acoustic tag through a midventral incision and were monitored for 92 days following the procedure. Instances of mortality and tag loss were recorded during this time. Necropsies were conducted to verify tag retention as well as to characterize the internal anatomical responses to the tag. A meta-analysis was used to determine the general effects of intraperitoneal tags on fish mortality and tag retention.

## Materials and Methods

### Animal care and holding conditions

Atlantic Salmon post-smolts (n =180; Body mass = 208.7 ± 3.0 g; Fork length = 269.7 ± 1.4 mm) were sourced from a commercial hatchery (Merlin Fish Farm Ltd., Wentworth, NS, Canada) in August of 2017. Fish were held as two shoals (n = 90 tank^-1^) in 3000 L circular tanks at the St. Andrews Biological Stations (St Andrews, New Brunswick, Canada). Tanks were maintained on a flow-through of filtered oceanic sea water (30 L min^-1^) at 12 °C under a 12 h dark: 12 h light photoperiod. Fish were fed *ad libitum* daily on a diet of commercial salmon feed (Optiline MB 200, Skretting Canada, Vancouver, British Columbia, Canada). Animals were held for two weeks to acclimate to lab conditions. All experimental procedures were conducted in accordance with guidelines established by the Canadian Council for Animal Care (CCAC) under approval of the Fisheries and Oceans Regional Animal Care Committee (RACC; AUP #17-04).

### Acoustic transmitter implantation procedures

This experiment assessed the impacts of the tagging procedure and tag type on Atlantic Salmon mortality and tag expulsion. Dummy tags provided by the manufacturer, made of the same material, weight, shape and size without electronic were used for implantation into the test fish. Salmon were haphazardly assigned to one of three treatment groups (n = 60 treatment^-1^): a sham control, or the fish was fitted with either a V7 tag (V7TP-2L; dimensions: 22 mm length x 3.5 mm radius; weight = 1.7 g in air; Innovasea Systems Inc., Bedford, Nova Scotia, Canada) or a V8 tag (V8-4L; 21 mm length x 4 mm radius; weight = 2 g in air). Each dummy tag had a unique serial number to help in identifying an individual fish. In the case of sham controls, animals underwent anaesthetization and the handling aspects of the surgical procedure, however, a tag was not implanted in the animal. Both the V7 and V8 tags were not actively transmitting throughout the experimental series (i.e., dummy tags). Each fish was fitted, intramuscularly, with a small Passive Integrated Transponder (PIT) tag while under anaesthetic to positively identify each fish.

Prior to surgical manipulations, all fish were fasted for at least 24 h. On the day of the tagging, individual fish were netted from the holding tank and immediately transferred to a bath containing Syncaine (aka tricaine methanesulfonate [MS222]; 100 mg L^-1^; Syndel; Nanaimo, BC, Canada) to sedate the animal. Once fish had been visibly sedated (i.e. lack of reactivity, low opercular rate), fork length and body mass were quickly taken, and the animal promptly transferred to a soft-foam V-trough. Restrained smolts had their gills continuously irrigated with aerated saltwater recirculated over their gills using a small hobby pump and a tube. This water contained a lower strength maintenance dose of Syncaine (50 mg L^-1^) to ensure that animals remained sedated throughout the entirety of the procedure. A small incision (∼20 mm) was then made on the ventral midline of the fish, and the tag gently inserted into the celomic cavity. The incision was then closed using up to three sutures (at 0.5-mm intervals) at the middle and ends of the wound (4-0 Vicryl sutures; Ethicon Inc., Raritan, New Jersey, USA). The fish recovered in a bath of raw saltwater until their righting reflex and opercular movements returned to normal. Recovered fish were then moved to and held in one of two large circular tanks (∼1000 L), each containing an equal mixture of fish that underwent one of the three treatments (n = 90 fish tank^-1^).

### Monitoring of chronic tag effects

Tagged fish were monitored over a total of 92 days and observed daily for any moralities or tag expulsions. The latter was determined based on acoustic tag IDs, and matched to an individual by PIT tag at the end of the experiment. However, not all instances of dummy tag expulsion were immediately recognized as they were occasionally consumed upon deposition in the tank. In such instances, tags were discovered in necropsies and thus had no exact time associated with expulsion. At the cessation of the monitoring period, each fish was euthanized using a lethal dose of Syncaine (150 mg L^-1^). Fish weight and length were taken for determinations of growth-related indices and length-weight relationships (see below). External abnormalities related to the tagging procedure were also characterized. This included if there was tag protrusion of the body wall, if sutures were missing, or if the incision had fully healed. Post-smolts were then dissected to confirm the presence of the tag within the coelom and to characterize how the internal structures had reacted to the tag. In the case of the former, this was noted as one of three outcomes: retained, expelled, or in progress of expulsion, meaning that the tag was encapsulated in the body wall of the fish. With respect to the reaction of the internal structures, this included characterization of any organs or tissues that the tag had become lodged in or incorporated into. Any signs of abnormalities, such as infection or tissue damage, were also noted. The fish’s sex was also determined during dissections.

### Calculations and statistical analyses

All statistical analyses were made using the R programming language (v 4.1.1) using R Studio (v 1.4.1717; [36]). All raw data for both the experimental and meta-analysis portions of this project and R scripts used in this project can be found here: https://github.com/mlaw27/Salmon-tag-retention-2023?search=1 . For all statistical models, significance was accepted at α = 0.05 with all values being mean ± SEM, unless otherwise noted. Survival analyses were conducted using the package ‘survival’ [37,38]. Specifically, we used a Cox proportional-hazards model to ascertain the effects of tagging treatment (i.e. sham, V7, or V8) on the time to morality and time to tag expulsion. In the case of the latter, sham fish were not included in the model as there was no tag to expel. It is worth noting that no loss of PIT tags was documented in any of the fish. In both models, fish that survived the entire experimental series (i.e. 92 days) or that did not expel their tags were censored from the analysis. Survival plots were made using the R package ‘survminer’ [39]. Comparisons of tag rejection status between V7 and V8 fish were made using a chi-square test of independence. Due to low sample sizes, the anatomical responses to each of the tags was not evaluated using statistical analyses and remain qualitative. Instantaneous growth rate (g) was determined as follows where Δt is the change in time in days, and w1 and w2 are the initial and final body masses of the fish (in grams), respectively [40].

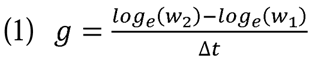

Specific growth rate (G) was calculated as follows using the fish’s instantaneous growth rate (g) [40]:

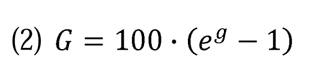

Specific growth rate was then analysed for tagging-related effects using a linear mixed model using the package ‘lme4’ [41], which included SGR as the response variable with the main effects of tagging treatment (i.e., sham, V7, or V8) and sex and the random effect of holding tank. We also characterized the relationship between the change in absolute body mass in relation to the absolute change in length as a measure of condition of the fish (i.e., a larger increase in body mass relative to a given change of length was indicated that the condition of the fish increased during the study period) using the same linear mixed model. This model also included the covariates tagging treatment and sex as main effects with holding tank as a random effect. In both models, pairwise comparisons were made between the tagging treatments (sham, V7, or V8) were made using a Tukey test [42] using the R package ‘emmeans’ [43]. To ensure that model assumptions were met, growth data were visually inspected for adherence to normality and equal variance using a Q-Q plot and a residuals vs. fit plot, respectively.

### Literature search and metanalysis procedures

Literatures searches were made using both Google Scholar (November 25 2021) and Web of Science (December 1 2022) with the following search terms: ((fish*) AND (“acoustic telemetry“ OR acoustic*) AND (tag shedding* OR tag expulsion* OR tag shedding* OR retention* OR expulsion* OR tag loss*) OR (tag* mortality OR survival OR death OR mortality)). For the Google Scholar results, we downloaded all publications from the first seven results pages constituting a total of 119 entries. For the Web of Science results, we selected the first 100 search results. Resulting lists were then first screened for their relevance by skimming the title and abstract to determine if it involved fish and tagging-based projects. From this preliminary list of papers, the specific details were extracted, which included the relevant citation information, taxa, species/common names, habitat (fresh, brackish, or saltwater), tag dimensions, type, and weight, incision location and size, and group sample sizes. We further refined this list to only include works that were using intracoelomic tags that did not extend outside the body cavity (e.g. radio tags with antennae) or gastric tags. Consequently, our results consisted of studies using intracoelomic dummy tags, acoustic tags, and passive integrated transponder (PIT) tags and were limited to only those reporting tag loss and/or tag-associated mortality. To determine tag to body mass ratio, reported values were used, if available. Otherwise, tag:body mass ratio was determined using the reported average body and tag mass. In instances where only a range of tag:body mass ratios are presented, we opted to extract the higher end of the range as a more conservative measure of this metric (i.e. if effects do exist, this should produce the largest effect size). Incision size always used the upper range of the size if this data was provided. Each experimental treatment was considered as an independent estimate, and in some cases, studies had multiple estimates stemming from either testing multiple tag types or having multiple experimental species. Each of these individual estimates was coded with a unique experimental number and nested within the paper ID to ensure that model estimate were not producing pseudoreplication (see below). Experimental duration consisted of the maximum monitoring duration of a particular trial, if indicated.

Statistical metanalysis procedures were conducted using the R package ‘metafor’ (V 3.4.0; [44]). Effect sizes and the corresponding variance for both mortality and number of tags retained were determined using the ‘escalc’ function. This involved expressing effect sizes as a proportional value, which were transformed using a Freeman-Tukey double-arcsine transform to meet model assumptions [45]. Transformed data was then analysed using a multivariate/multilevel linear (mixed-effects) model fit using a restricted maximum likelihood approach (REML; [46]). For both tag retention and mortality estimates, we treated our models in a stepwise fashion by first assessing a complete model with all fixed effects (tag length, tag diameter, incision size, experiment duration, and tag:body mass ratio) and then assessing each fixed effect individually against the response variable. This was done as some articles did not include all of the fixed-effects metrics and thus may not have been represented in the full model. To ensure that we accurately portrayed these metrics on affecting response variables, we opted to use the individual models as well. For all models, we also included paper ID as a random effect, which had a nested term of experimental number included for each work thereby accounting for any repeated sampling that may have occurred. *P* values for all model terms were also corrected for false discovery rates using a Benjamini-Hochberg correction [47]. All model estimates/outputs are presented in the transformed data. Models were visually inspected using profile likelihood plots [44,48].

## Results

### Post-tagging mortality

Mortality associated with the surgical procedures and implantation of acoustic tags was minimal across all treatment groups (≤ 5.1%; Fig. 1). Interestingly, of the six post-smolts that died, two appeared to be quite thin and small suggesting some underlying physiological/anatomical issues. The remainder of these fish did not have any other obvious symptoms or signs that may have explained their mortality.

**Figure 1:**
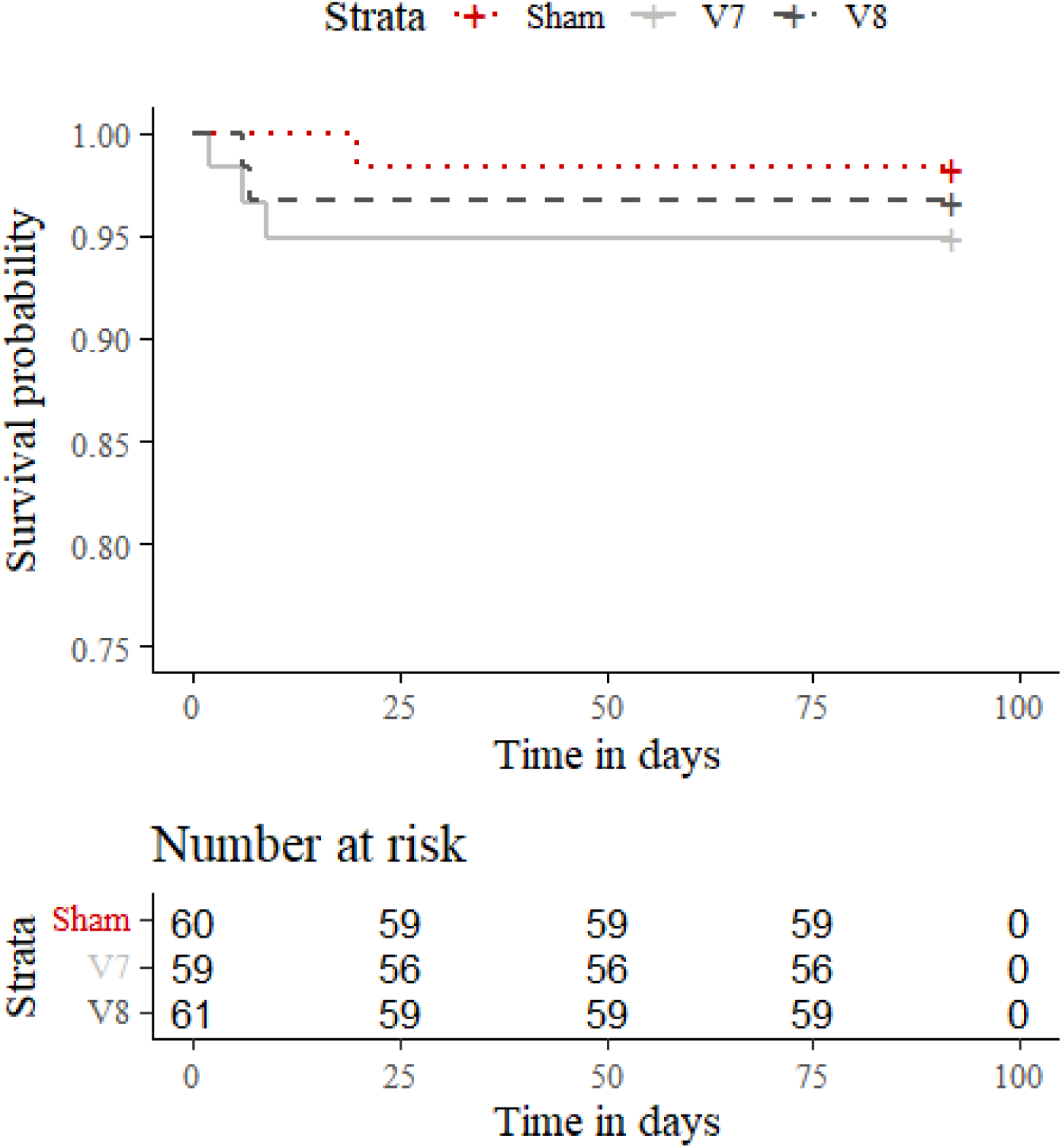
Plot representing the survival of Atlantic salmon smolts following the a sham surgery (red, dotted line), or the intracelomic insertion of a Innovasea V7 (black, dashed line) or V8 (grey, solid line) acoustic tag. Fish were monitored over a 92-day time frame with survivors censored from the analysis (denoted by a ‘+’ shape).

### Tag retention

Rates of tag retention were fairly poor among Atlantic salmon post-smolts. Overall, 54.8% of fish had either shed their tag already or were in the process of expelling their tag over the 92-day monitoring period. A total of 34.8% percent of all tagged fish had their tag fully expelled from the body cavity. Of those that fully expelled their tag and that had a recorded expulsion date, the overall mean time to tag expulsion was 33.3 ± 1.1 days (n = 20). On a tag-treatment basis, V7 and V8 transmitters had comparable times to tag expulsions (Likelihood ratio test= 1; df = 1; *P* = 0.3; Fig. 2) suggesting that transmitter size may not be a factor of importance. This is further supported by comparisons of the observed status of the tags at the termination of the experiment (i.e. retained, expelled, or in progress) being similar between fish fitted with V7 and V8 transmitters (χ^2^ = 1.99; df = 2; *P* = 0.4; Fig. 2). More specifically, only 48% and 42% of post-smolts fully retained their tags by the end of the 92-day monitoring period for V7-and V8-fitted fish, respectively. Although, tagging group had no effect on tag expulsion, it did appear that V8-fitted fish had a greater percent of fully expelled tags when compared to V7-fitted fish (40.7% vs. 28.6%).

**Figure 2:**
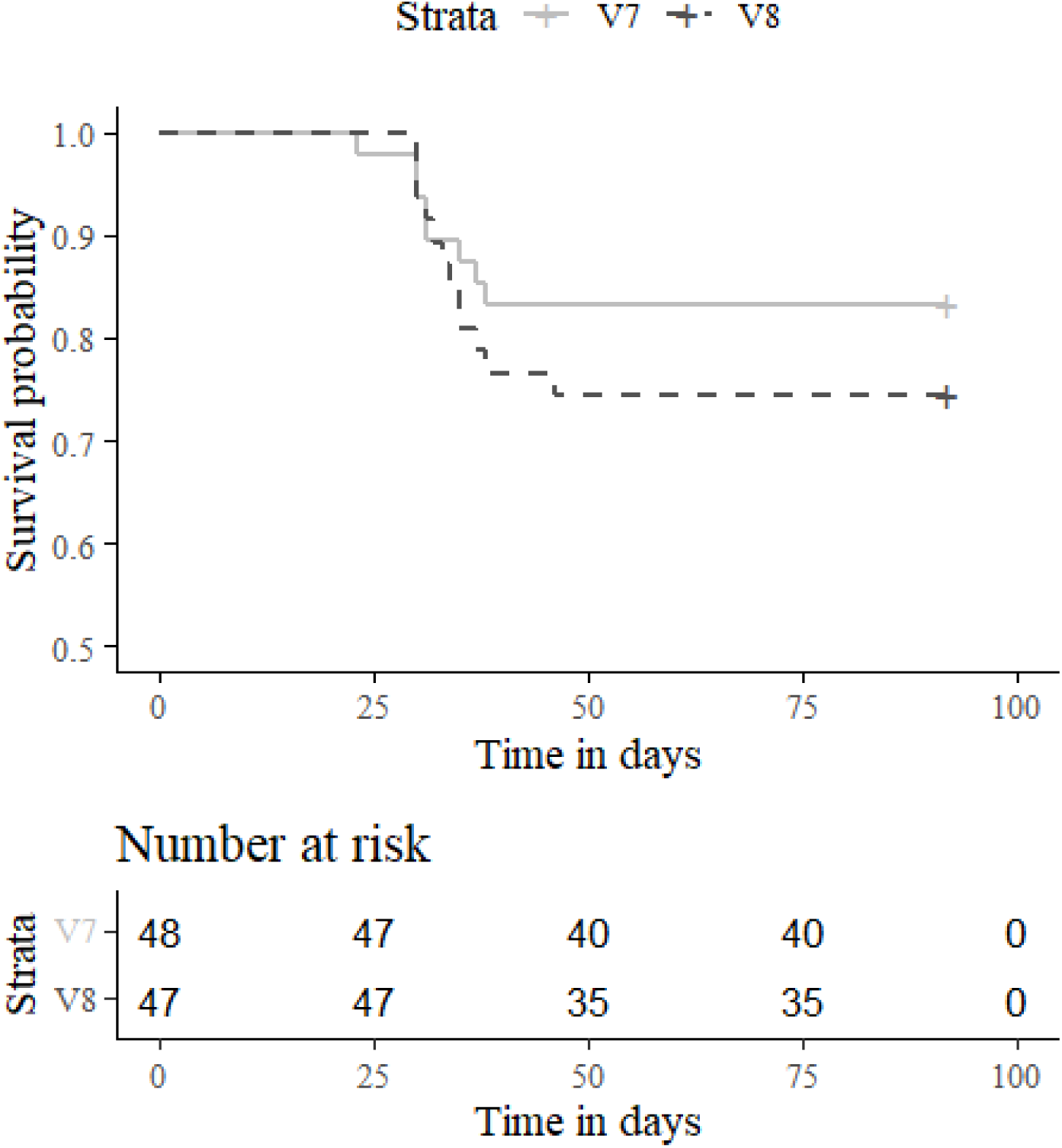
Plot representing the number of days that a intracelomic acoustic tag was retained inside the body cavity of an Atlantic salmon post-smolt following the insertion of either a Innovasea V7 (black, dashed line) or V8 (grey, solid line) acoustic tag. Fish were monitored over a 92-day time frame for tag expulsion through daily checks, with fish retaining the tag until the end of the experiment censored from the analysis (denoted by a ‘+’ shape).

Figure 3 exemplifies the differences between a tag that was retained and a tag in progress of expulsion. In fish that retained their tag, there was no external signs of the tag being forced from the body (Fig. 3A). Alternatively, the start of expulsion appeared to result from tag encapsulation by body wall mesentery (Fig. 3B) and then becoming lodged in the dorsal musculature (Fig. 3C). Typically, this was away from the incision site and occurred on the lateral surface of the fish. From an external view, an ‘in progress’ tag expulsion demonstrated the slight bulge in the skin (Fig. 3D) with gradual thinning and eventual rupture through the skin into the external environment (Fig. 3E). Fish that retained the tag completely often had the tag encapsulated by the mesentery (Fig. 3B) or was lodged into one of the internal organs, such as the pyloric caeca (Fig. 3F).

**Figure 3:**
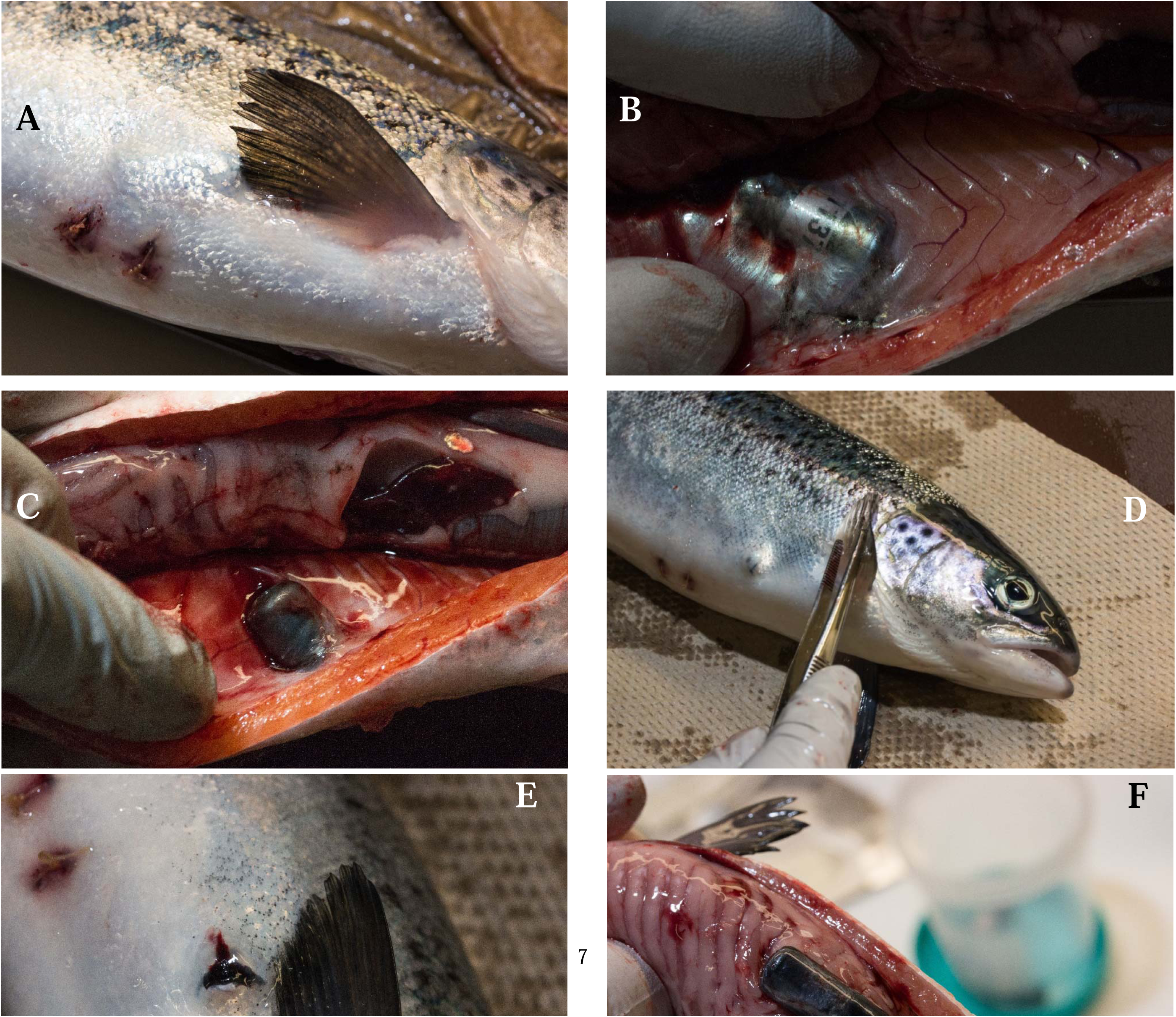
Images depicting the fates of acoustic tags in post-smolt Atlantic Salmon following a 92-day monitoring period. In image **A**, the incision from tag insertion (with retained sutures) has completely healed and there is no external signs of tag rejection. An example of tag has been encapsulated by mesentery in the coelom of the fish (**B**). The tag is starting to be expelled through the lateral surface of the dorsal musculature away from the incision (**C**). The expulsion of the tag can be seen first as a bulge on the lateral surface of the fish (**D**), culminating in it rupturing through the skin (**E**). In the case of fish that did retain their tags, the tag was often seen lodged in one of the organ structures such as in the pyloric ceca as depicted here (**F**).

We also quantified the fate of the tags within the body cavity of the fish. While no statistical analyses were ran, most of the tags that were retained within the body cavity were encapsulated by the mesentery for both treatment groups (Fig. 4). These retained tags were largely concentrated to the right lateral and medial, of the left-right and antero-posterior axes, respectively. The other most common response in fish that retained tags was that the tag was not encapsulated by any internal organs or structures (Fig. 4). In V7-fitted fish, one tag was found to be embedded in the pyloric caeca while a singular fish with a V8 tag had the surrounding tissues adhering around the tag (Fig. 4).

**Figure 4:**
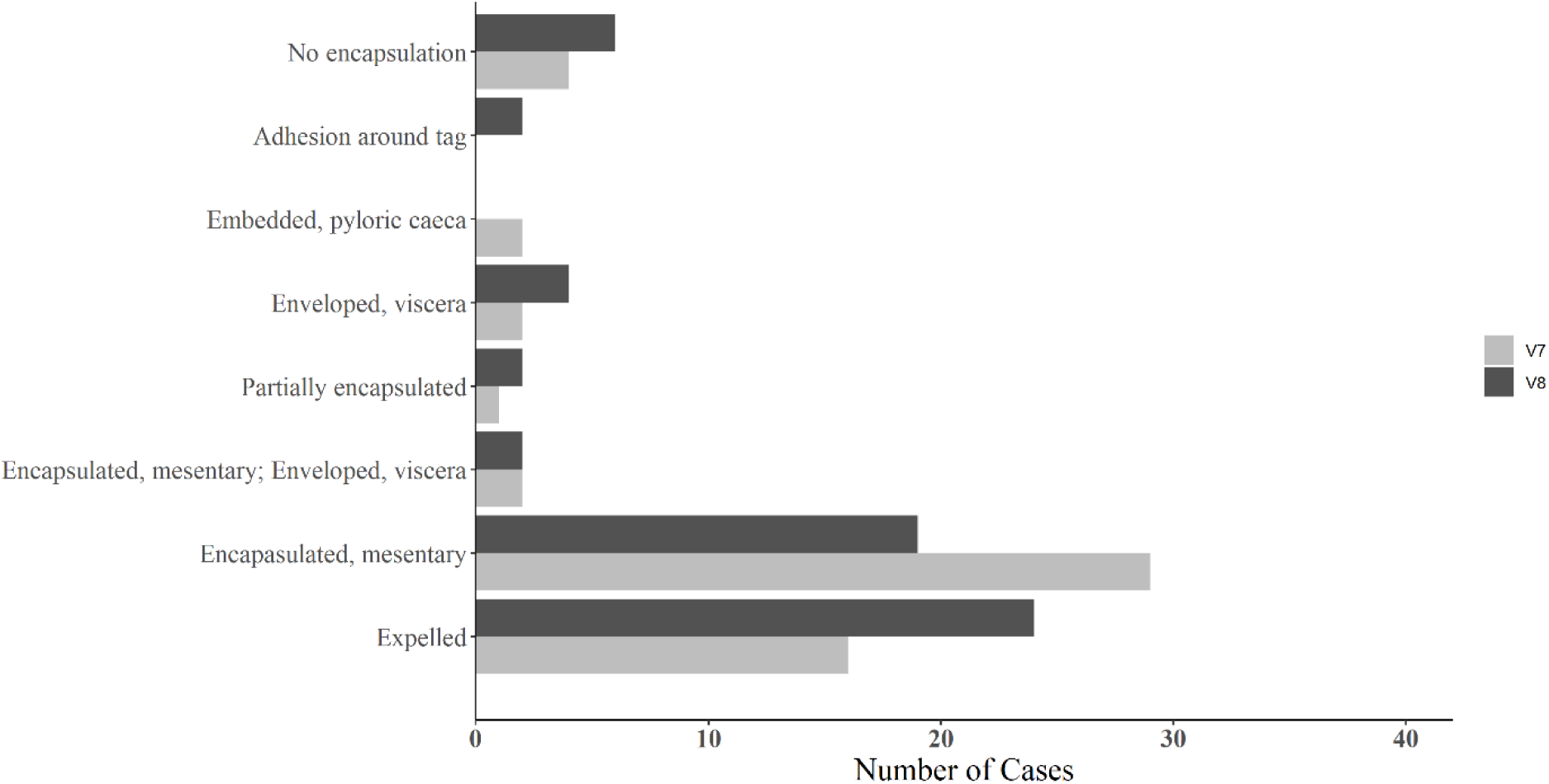
Counts of the final anatomical location of an intracelomic tag after 92 days being fit within an Atlantic salmon smolt. Smolts were fitted with either a Innovasea V7 (light gray) or a V8 (dark grey) acoustic transmitter. These two treatment groups were not statistically compared against one another due to low sample sizes.

### Growth and condition outcomes

Tagging treatments impacted the growth indices investigated. There was a significant, positive relationship between the change in a fish’s body weight and the change in the fish’s fork length (df = 166.48; t = 18.45; P < 0.001) such that individuals that had larger changes in body mass also had larger length changes (Fig. 5). There was no effect of either tagging type or sex on this relationship (Table 1). The use of the V7 tags resulted in lower SGR (*P* =0.002; Fig. 6) compared to sham controls, being 13.47% lower than for sham fish (Table 1). In contrast, fish tagged with V8 transmitters were comparable to both shams and the V7 tagged fish (Fig. 6; Table 1).

**Figure 5:**
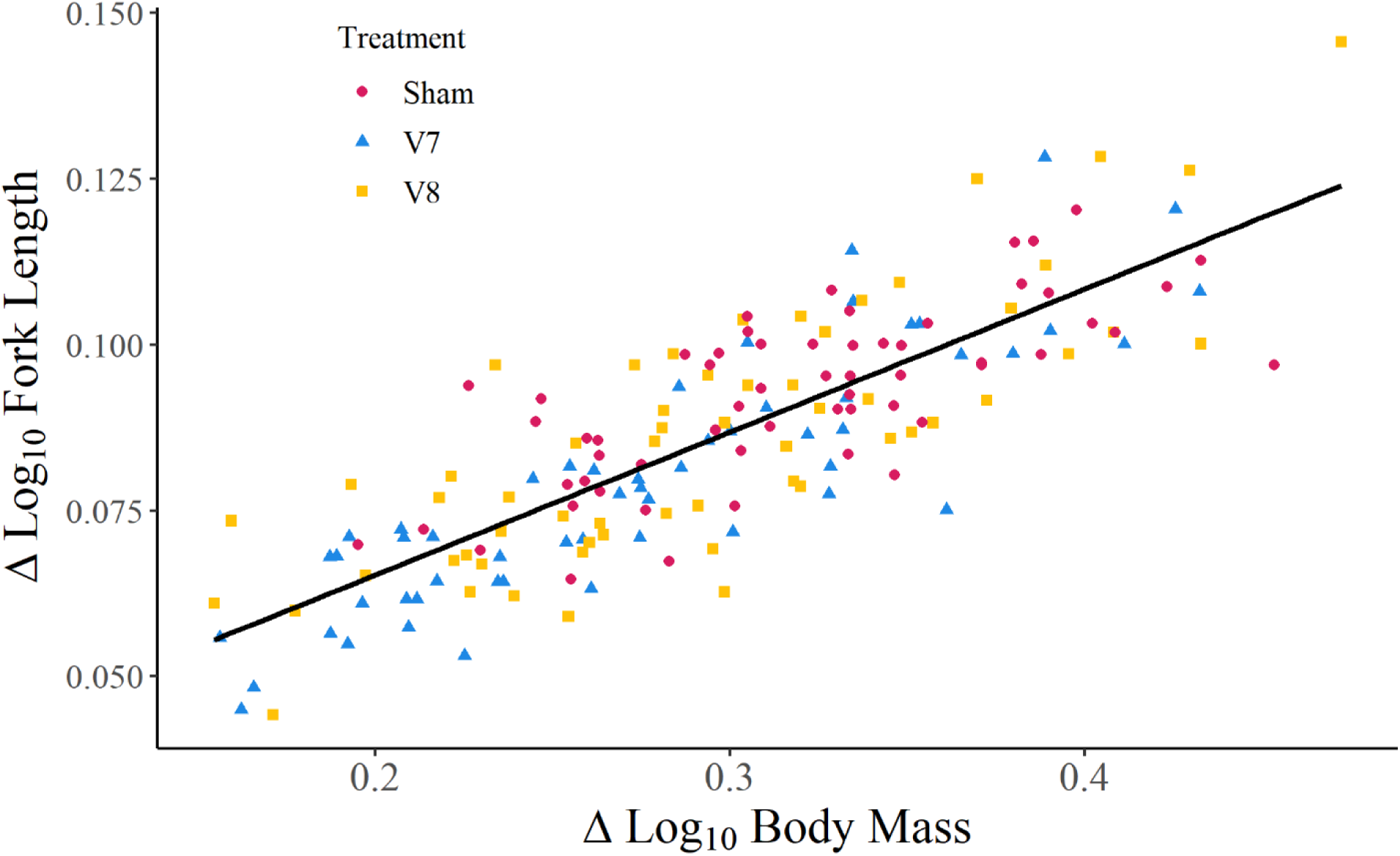
Relationship between an individual’s change in Log_10_ body mass with its change in Log_10_ fork length over a period of 92 days. Animals were treated as either a surgical sham or implanted with an Innovasea V7 or V8 acoustic tag. A linear mixed effects model was fit to the data with the change in Log_10_ fork length as a product of the change in Log_10_ body mass, tagging treatment, and the fish’s sex as the main effects and tank as a random effect. Statistical significance was accepted at α = 0.05. Changes in Log_10_ fork length were found to be the product of changes in Log_10_ body mass (*P* <0.001) whereas both tagging treatment and sex had no effect. The black line represents the overall relationship of the change in Log_10_ body mass with its change in Log_10_ fork length independent of tagging or sex effects.

**Figure 6:**
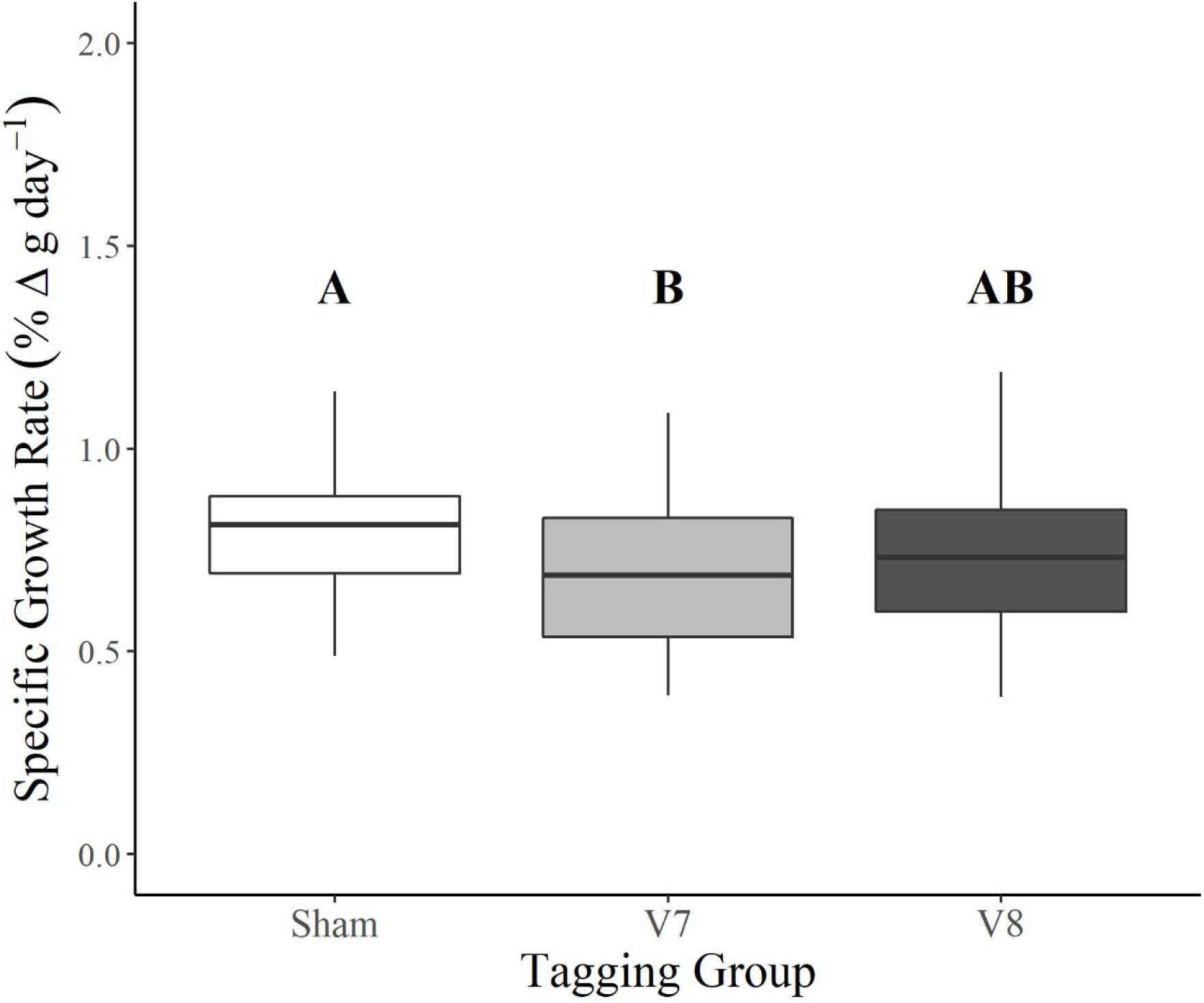
Effects of a sham surgery (white) and the insertion of either a Innovasea V7 (light grey) or a V8 (dark grey) acoustic tag on the specific growth rate (SGR). Statistical significance was accepted at α = 0.05. A linear mixed effects model was fit to the data and, where main effects were significant, a Tukey test was used to discern pairwise differences among treatment groups. The fish’s sex was also included as a fixed effect in the model and tank was a random effect.

**Table 1:**
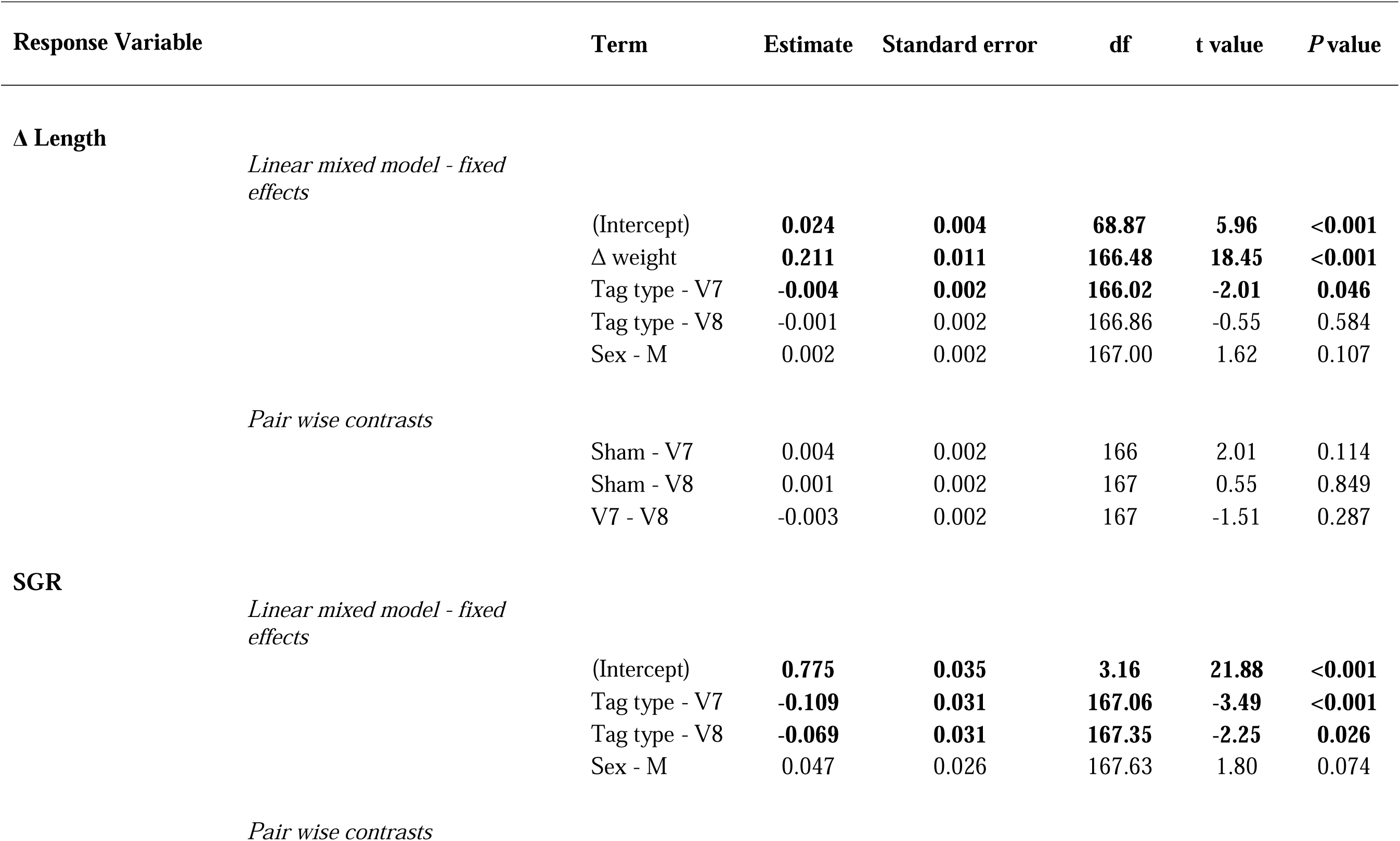
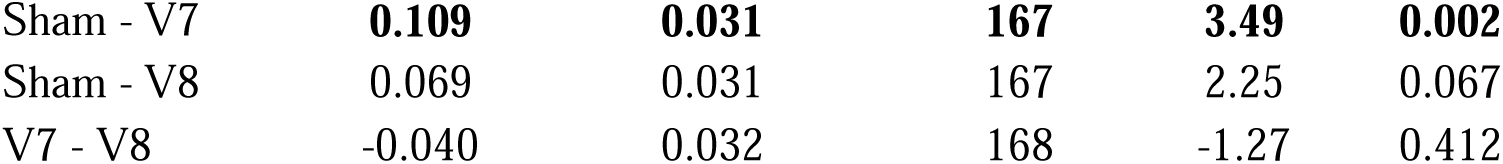
Statistical output for the effect of tag type (i.e. sham control, V7 or V8 acoustic tags) and sex on the condition factor and growth indices in Atlantic salmon smolts monitored over a 92-day holding period. A linear mixed model with the fixed effects of Tag type, sex, and Δ weight (Δ length model only) and a random effect of holding tank was used to analyse the response variable. Tag type effects were addressed using Tukey pairwise contrasts. Bolded rows represent statistically significant results (α = 0.05).

### Metanalysis summary and metaregression statistics

Collective mean mortality rate was 12.4% and ranged between 0% and 90%. In the full model, tag diameter was the only fixed effect found to influence tagging mortality across our studies and was positively associated with it (Table 2). However, when looking at just tag diameter against mortality in the individual models, there was no relationship between the two parameters (Estimate = 0.044; 95% CI_L_ = -0.007, 95% CI_u_ = 0.021; Adjusted *P* = 0.346; Table 2). None of the other fixed effect parameters exhibited statistically significant relationships with mortality in the individual models (Table 2).

**Table 2:**
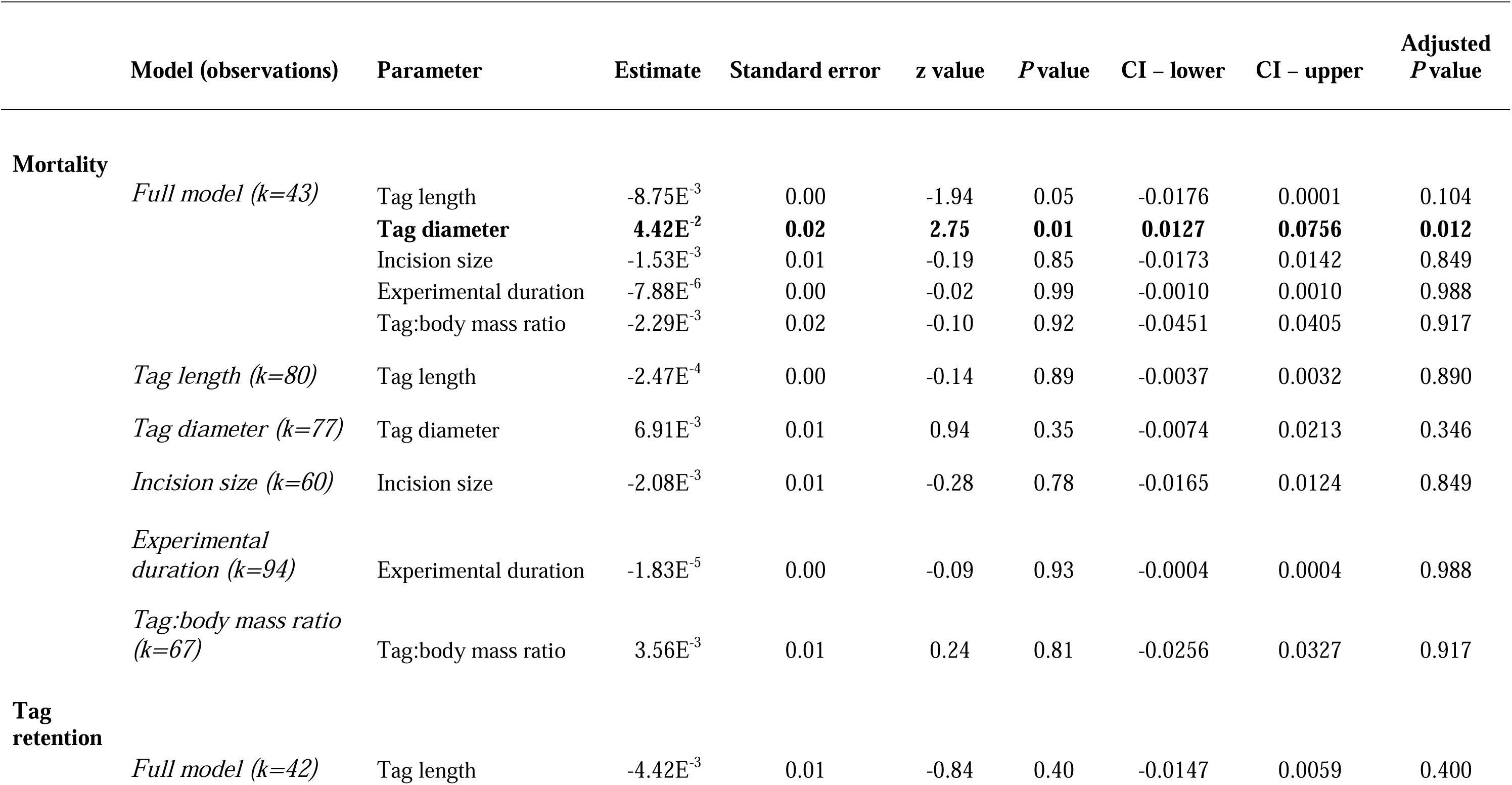
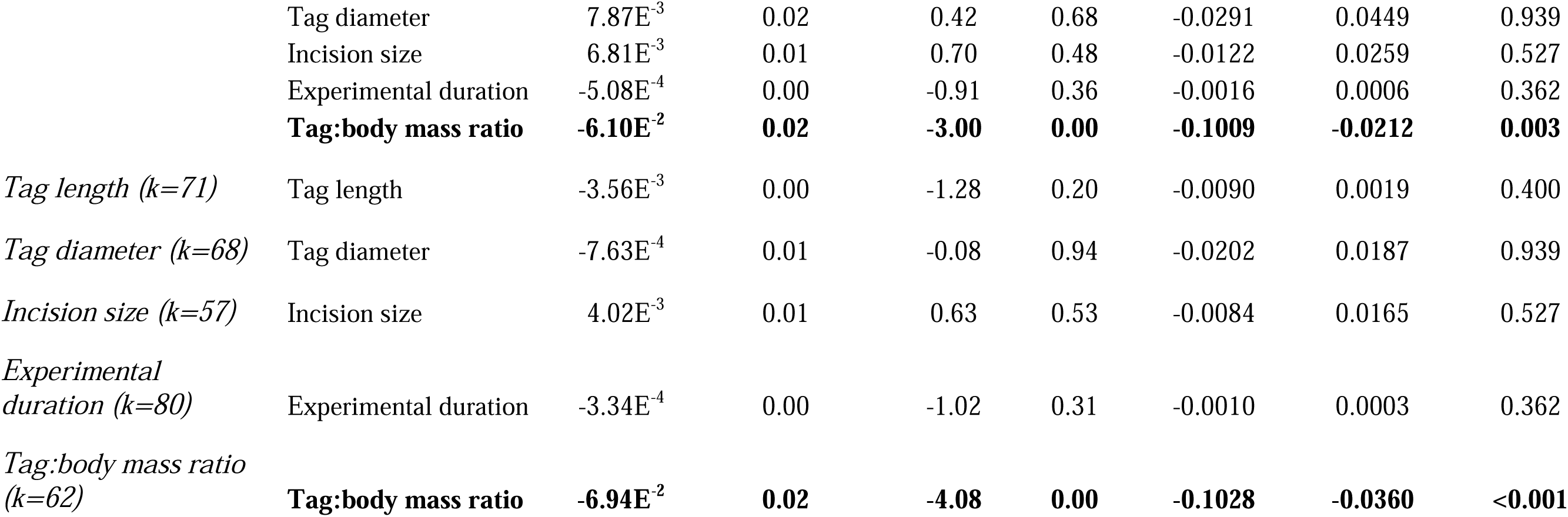
Meta-regression fixed effects model estimates comparing tagging and experimental parameters against tag retention and mortality rates in fishes. The term ‘Full model’ represents a meta-regression containing all fixed effects while subsequent models examined a single fixed effect against the response variable. Meta-regressions were conducted using proportional data transformed using a Freeman–Tukey double-arcsine transformation. Statistical significance was accepted at α = 0.05 with P values being corrected for multiple comparisons using a Benjamini-Hochberg correction. Significant terms are bolded in the table. k values represent the number of observations for a specific model.

|  | Model (observations) | Parameter | Estimate | Standard error | z value | P value | CI – lower | CI – upper | Adjusted P value |
| --- | --- | --- | --- | --- | --- | --- | --- | --- | --- |
| **Mortality** | *Full model (k=43)* | Tag length | -8.75E<sup>-3</sup> | 0.00 | -1.94 | 0.05 | -0.0176 | 0.0001 | 0.104 |
|  |  | **Tag diameter** | **4.42E<sup>-2</sup>** | **0.02** | **2.75** | **0.01** | **0.0127** | **0.0756** | **0.012** |
|  |  | Incision size | -1.53E<sup>-3</sup> | 0.01 | -0.19 | 0.85 | -0.0173 | 0.0142 | 0.849 |
|  |  | Experimental duration | -7.88E<sup>-6</sup> | 0.00 | -0.02 | 0.99 | -0.0010 | 0.0010 | 0.988 |
|  |  | Tag:body mass ratio | -2.29E<sup>-3</sup> | 0.02 | -0.10 | 0.92 | -0.0451 | 0.0405 | 0.917 |
|  | *Tag length (k=80)* | Tag length | -2.47E<sup>-4</sup> | 0.00 | -0.14 | 0.89 | -0.0037 | 0.0032 | 0.890 |
|  | *Tag diameter (k=77)* | Tag diameter | 6.91E<sup>-3</sup> | 0.01 | 0.94 | 0.35 | -0.0074 | 0.0213 | 0.346 |
|  | *Incision size (k=60)* | Incision size | -2.08E<sup>-3</sup> | 0.01 | -0.28 | 0.78 | -0.0165 | 0.0124 | 0.849 |
|  | *Experimental duration (k=94)* | Experimental duration | -1.83E<sup>-5</sup> | 0.00 | -0.09 | 0.93 | -0.0004 | 0.0004 | 0.988 |
|  | *Tag:body mass ratio (k=67)* | Tag:body mass ratio | 3.56E<sup>-3</sup> | 0.01 | 0.24 | 0.81 | -0.0256 | 0.0327 | 0.917 |
| **Tag retention** | *Full model (k=42)* | Tag length | -4.42E<sup>-3</sup> | 0.01 | -0.84 | 0.40 | -0.0147 | 0.0059 | 0.400 |

|  |  |  |  |  |  |  |  |  |
| --- | --- | --- | --- | --- | --- | --- | --- | --- |
|  | Tag diameter | 7.87E<sup>-3</sup> | 0.02 | 0.42 | 0.68 | -0.0291 | 0.0449 | 0.939 |
|  | Incision size | 6.81E<sup>-3</sup> | 0.01 | 0.70 | 0.48 | -0.0122 | 0.0259 | 0.527 |
|  | Experimental duration | -5.08E<sup>-4</sup> | 0.00 | -0.91 | 0.36 | -0.0016 | 0.0006 | 0.362 |
|  | **Tag:body mass ratio** | **-6.10E<sup>-2</sup>** | **0.02** | **-3.00** | **0.00** | **-0.1009** | **-0.0212** | **0.003** |
| *Tag length (k=71)* | Tag length | -3.56E<sup>-3</sup> | 0.00 | -1.28 | 0.20 | -0.0090 | 0.0019 | 0.400 |
| *Tag diameter (k=68)* | Tag diameter | -7.63E<sup>-4</sup> | 0.01 | -0.08 | 0.94 | -0.0202 | 0.0187 | 0.939 |
| *Incision size (k=57)* | Incision size | 4.02E<sup>-3</sup> | 0.01 | 0.63 | 0.53 | -0.0084 | 0.0165 | 0.527 |
| *Experimental duration (k=80)* | Experimental duration | -3.34E<sup>-4</sup> | 0.00 | -1.02 | 0.31 | -0.0010 | 0.0003 | 0.362 |
| *Tag:body mass ratio (k=62)* | **Tag:body mass ratio** | **-6.94E<sup>-2</sup>** | **0.02** | **-4.08** | **0.00** | **-0.1028** | **-0.0360** | **<0.001** |

Across all studies, tag retention in fishes averaged 86.7%, ranging from 20% to 100% retention. In the full model, tagging retention was only affected by tag:body mass ratios (Estimate = -0.061; 95% CI_L_ = -0.101, 95% CI_u_ = -0.021; *P* = 0.003; Table 2) such that increasing tag:body mass ratios are associated with a decrease in the likelihood of tag retention (Fig. 7). No other fixed effect had a statistically significant effect on tag retention in the full model, which was a pattern also shared by the individual models (Table 3). Point estimates and the corresponding confidence intervals as well as the models’ meta-regression variance components for all the tag retention and mortality values used here can be found in the supplementary materials.

**Figure 7:**
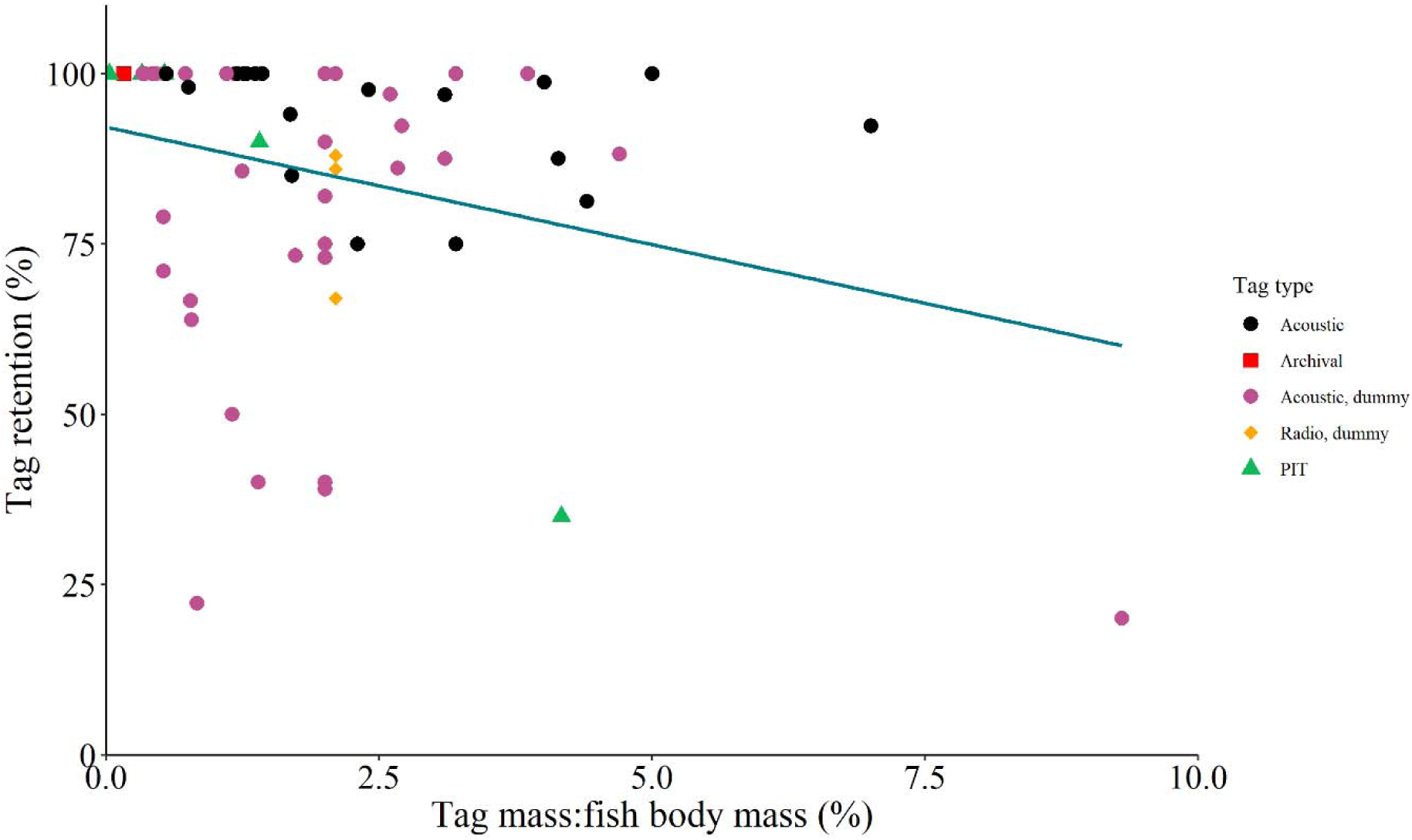
Scatter plot depicting the relationship between an intracoelomic tag being retained and the tag to body mass ratio. In the metaregression, this relationship was statistically significant (P < 0.05) and is visually denoted by the blue line. Each point represents a unique experiment’s tag retention rate across multiple studies and the type of tag is denoted by both colour and shape.

## Discussion

### Overview

This study attempted to characterise rates of mortality and tag retention in Atlantic salmon post-smolts fitted with a dummy acoustic tag. Secondarily, we conducted a formal metanalysis of the literature to assay covariates that may influence tag loss and post-tagging mortality to discern areas where tagging procedures could be refined to minimize tag losses or mortality. Mortality in our salmon was consistently low across treatment groups and was consistent with the literature at large. However, both the results of our experiment and of the meta-analysis are unable to address specific causal mechanisms underlying post-tagging mortality and suggest that context-specific effects, especially stress status, are likely the main driver of mortality. With respect to tag retention, rates of tag loss were high in this study, in contrast to general findings in the literature, and may stem from a heightened foreign body response. Growth of tagged salmon post-smolts was also mildly impaired and may suggest increased metabolic loading associated with a tagged state. Together, we recommend that tagging be conducted in a manner that minimizes stress and that uses tag sizes that are appropriately scaled to the focal fish of interest (i.e., as small as possible). These factors would likely minimize tag-related mortality while increasing tag retention to ensure that experimental series are reflective of fish behaviours and not misinterpreted as additional mortality (i.e., tag expelled, but fish survived).

### The effects of tagging in modulating mortality rates

Tag implantation appeared to have little impact on mortality of post-smolt salmon. This result is consistent with the literature, where the use of intraperitoneal tags is often associated with low rates of mortality (<10%) in salmonids [21,26,49–51] and, more broadly, teleost fishes [52–54]. Similarly, our metanalysis concluded that mean post-release mortality rate was 12.4% across all studies examined, suggesting that the procedures used to implant tags appeared to have a minimal impact on post-release survival, all else being equal. However, caution must be exercised in this statement as tagging-mortality studies are often conducted in a controlled, laboratory-based environment where the fish can recover under ideal environmental conditions and in the absence of predators [27]. It is entirely possible that the additional stress burden imparted by surgical procedures may increase susceptibility to mortality in the wild, especially if a fish is facing several energetically-demanding processes simultaneously (e.g. temperature shifts, infection, sustained swimming, predator evasion). For Atlantic salmon post-smolts, tagging-associated mortality is expected to be quite low based on our findings.

Tagging-mortality rates are also not necessarily low across all settings and can be highly context-specific. While the experimental portion of this study demonstrated low tagging-associated mortality, there was considerable variation in study-specific mortality rates in the meta-analysis, ranging from 0 to 90 % mortality. Several factors can exacerbate post-release mortality of tagged fish, including: fish body mass and length [49,59–61], tag size [51,59,,61], tag to body mass ratio [31,32,62,63], and the source population of fish [51]. Based on our meta-regression results, these factors appear to have little influence on mortality, suggesting that responses are either highly context-specific (i.e. mortality arises under a specific set of conditions or in a specific species) or that there is largely a null effect. For example, the ‘2% rule’ (i.e. tag mass remains under 2% of the fish’s body mass [64]) was commonly believed to be an important consideration for minimizing tagging mortality and behavioural impairments in fishes [29,65]. New evidence suggests that this effect is not ubiquitous; rather that contextual considerations should drive the appropriate tag size for the study in question [27,66,,67].

Similarly, the general lack of effect of the model covariates in our meta-regression reflects an analogous situation where context-specific effects of the experimental design are likely more important driving factors for tagging-related mortality. Together, our results suggest that there are no clear patterns related to tag parameters, incision size, or experimental duration affecting tagging-related mortality.

Despite the increasing popularity of acoustic tagging as a tool in fisheries science, we still lack a thorough understanding of the specific physiological mechanisms driving tagging-related mortality in fishes. Regardless of the specific methodology used, the process of tagging is generally considered to be stressful to fish across several levels. Capture [68–71], handling [72–74], and surgery/sedation [75–78] can all induce pronounced stress responses in fishes, which is often marked by elevations in metabolic rate and circulating levels of high energy substrates and cortisol as well as the development of acidosis, among a multitude of other effects. Indeed, stress biomarkers have been observed in fishes following the tagging procedure [79,80], although reporting on tagging-specific response are scant in the literature. At this time, we can only speculate that physiological perturbations associated with tagging are likely the main factor driving post-release mortality in tagged fish. Thus, we recommend that tagging be conducted in a manner that minimizes stress to the animal (i.e. minimize air exposure and handling, appropriate collection methods, minimizing captivity durations, ideal anaesthesia dosages [27,81–83]) and that pilot trials be conducted to ensure that tagging conditions are optimized for fish welfare (i.e. selecting appropriate tag size, tag insertion location, and recovery periods [27,84–86]). While there remains uncertainty regarding the physiological drivers surrounding post-tagging mortality, ensuring that stress is minimized should help enhance survival of tagged fish.

### Possible casual factors driving tag expulsion in fishes

The immune system likely plays a key role in determining tag retention rates in fishes. While our understanding of how the immune system mediates the coordination, isolation, and expulsion of tags in fishes appears rather limited [87–90], there is a wealth of knowledge concerning the foreign body response in a clinical setting [91,92]. Briefly, the response to a foreign object is coordinated by the immune system. The initial, acute phase of the foreign body response involves the release of proinflammatory factors that attracts neutrophils to the area, which further releases proinflammatory agents and increases localized vascularization. The neutrophils’ actions also attract monocytes to the area, which undergo differentiation to macrophages. The macrophages are the ‘workhorses’ of this system as they are responsible for enveloping the object and secreting degradatory compounds to eliminate the object while also further mediating the inflammatory response [91,92]. However, if the object cannot be broken down, the actions of the macrophages switches to a chronic, fibrosis-generating role by activating fibroblasts to produce a proteinaceous, extracellular matrix (ECM) to encapsulate and isolate the foreign body from the surrounding tissues [91,92]. This is proceeded by vascularization of the capsule and an increased proliferation of the ECM until a steady state of growth is achieved. In the mammalian model, this process appears to largely result in the foreign body being retained and isolated as in the case of various biomaterials and implants [91–94] but in some rarer instances, these objects can be expelled from the body [95,96].

In fishes, there is evidence that the foreign body response is important in mediating tag expulsion. One of the first works to address a causal mechanism underlying tag explosion found that in channel catfish (*Ictalurus punctatus*), like mammals, dummy acoustic tags were encapsulated in a layer of myofibroblasts and collagen tissue prior to expulsion [90]. Marty and Summerfelt’s [90] proposed model of expulsion was similar to mammalian models at the time albeit with expulsion driven by the capsule’s myofibrolasts contracting the tag against the body wall. Lucas [97] further elaborates on this model by suggesting that trans-body wall passage of encapsulated tags occurs through pressure necrosis at the exit point, which is supported by their histological characterisations therein. While our work did not address tissue histology specifically, we did note that tags were often encapsulated by fibrous tissue in the body cavity of the fish and that expelled tags were encapsulated and superficially appeared to move through a site experiencing localized tissue necrosis (see Fig. 6) akin to what Lucas [97] had previously described. The greater body of literature also appears to support a role of the foreign body response in mediating tag retention/expulsion, as evidenced by tag encapsulation or tags translocating across body walls/organ structures [87–89,98,99]. While some of our salmon did have clear signs of body wall expulsion, we cannot rule out expulsion via the anus given that transintestinal movement of tags can occur in fishes [90]. Together, our results provide further evidence in support of the foreign body response in mediating tag expulsion.

The percentage of tags that were expelled or in the process of expelling in our salmon was quite high. The 45% retention rate in our salmon was unexpected as prior work with salmonids demonstrates high retention rates (>85% [31,63,100–104]), which is further supported by our metanalysis results of the wider fish literature (∼87% retention). From an analytical perspective, low tag retention is problematic in a telemetry study as it may result in false positives of stationary tags and inferred mortality when in fact the tag has been simply discarded and surviving fish remains active. Consequently, this high rate of tag shedding should be accounted for in acoustic analyses when conducting trials with Atlantic salmon post-smolts. As with other works [32,49,,105], we found that when shedding occurred, it was early in the monitoring period (∼33 days) suggesting that losses are likely greater in the early stages following release. While we are unsure of the exact mechanism driving this effect in our salmon, tag expulsion rates can be modulated by several factors, including water temperature [87,102,,106], large/heavy tag size [32,90], and decreasing fish body size [49,63,102,105]. Our metanalysis results also supported this to an extent whereby we found a significant relationship between tag retention rates and tag-to-body mass ratios. However, we are sceptical that any of these factors are playing a role in the high rates of expulsion given that tag weights represented a small fraction of the fish’s body mass (∼1%), are a typical size for fish of this size class, and were reared in a ‘normal’ temperature range for this species [107]. In the clinical literature, the size, shape, location, and sterility of the object as well as any localized tissue damage can influence the magnitude of the foreign body response [92,94,,108]. While entirely speculative, the area that the tag settled in as well as localized bacteria/tissue damage prompted a heightened immune reaction in response to the tag. Future works should address the role of the foreign body response in mediating tag retention dynamics in fishes as it could provide valuable insight in developing predictive models of tag expulsion likelihood in a field setting.

The results of the meta-analysis provided a great deal of insight to assess important factors related to enhancing tag retention in fishes. While tag dimensions and mass were shown to not affect tag retention rates in the meta-regression, tag-to-body mass ratio did positively scale with tag expulsion in the greater literature, indicating this as a key consideration in designing telemetry studies. Rather than a hard cut-off as proposed in the 2% rule [64], it suggests that this relationship exists along a continuum where generally larger ratios correspond with a greater likelihood of expulsion, particularly at extreme tag:body weight ratios. In the context of the foreign body response, such comparatively large objects are likely to cause a greater immune response [92,94,,108] and may result in earlier expulsion [90]. Our results also indicate that the absolute dimensions of the tag (length and mass) are not nearly important as the relative size of the tag to the size of the fish. Consequently, we would recommend using the smallest possible tag size that also meets the experimental goals of the project (i.e., maximizing the trade-off in tag size to battery life) to ensure a minimal loss from expulsion. Of additional interest is that monitoring duration did not affect tag expulsion rates in the present study. In some works, most tag loss appears to occur over the first few weeks of the monitoring period with fewer losses happening over more chronic durations [32,49,,105]. While this may be a trend in the literature, it is important to understand that our metric of time in the meta-regression represents an average tag loss over the monitoring duration rather than time to tag loss as this latter value was often not reported. As such, it is possible that tag loss may be greater in the initial weeks following implantation and should be considered when designing an experimental series.

### Tag effects on growth rate

A key consideration when designing a telemetry study is avoiding adverse impacts of the tag itself on the behaviour and fitness of monitored fish [67,109]. Effects on growth parameters are often used as a proxy for tag-related impacts on the fish and have been widely explored in tagged fishes [24,26,33,49,97,110–112]. In the salmon investigated here, we found that tagging had a negative impact on SGR (13.47% lower for V7 tagged fish over a 92 day period), consistent with some of the literature [49,110,,112]. Likely, tag burden or a stress effect related to tag implantation is imparting a greater metabolic load on the animal resulting in reduced growth inputs. However, additional metabolic profiling would be needed to confirm such notions.

Furthermore, tagging-related impacts on growth appear to be highly contextual and can have a strong temporal component. For example, Greenstreet and Morgan [110] found that while small size classes of salmon parr (<160 mm) had lost body weight while larger conspecifics were still accruing body mass. Similarly, condition factor (K) of bloaters (*Coregonus hoyiinitially*) decreased following tagging but recovered in the latter half of the experiment. As our monitoring of growth was based on initial and final changes in mass/length values, we may not be addressing some of the more nuanced changes in growth rates here. Overall, these suggests do suggest that tag burden is associated with impaired growth in post-smolt Atlantic salmon.

## Funding

This study was supported from funding awarded by the Aquaculture Ecosystem Interactions Program at Fisheries and Oceans Canada.

## Acknowledgments

The authors thank the Atlantic Salmon Federation for kindly supplying the dummy tags that were used as part of the experiments described in this study.

---

**Table S1:**
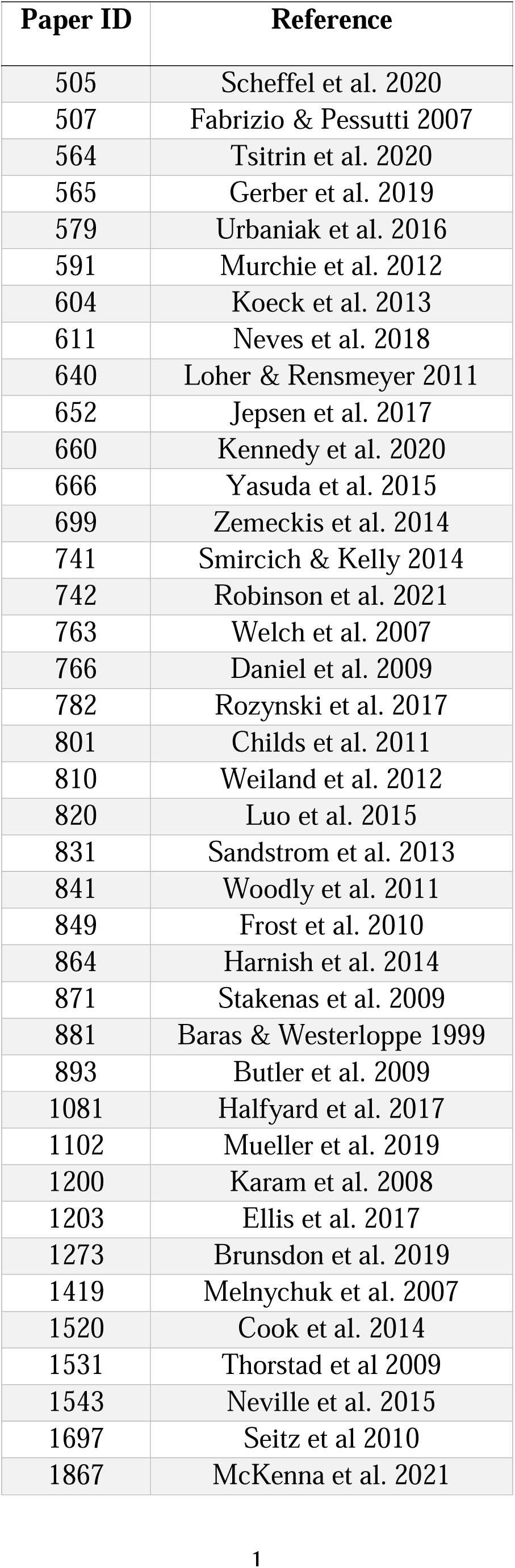
Paper identification key for the references used in the meta regression.

**Table S2:**
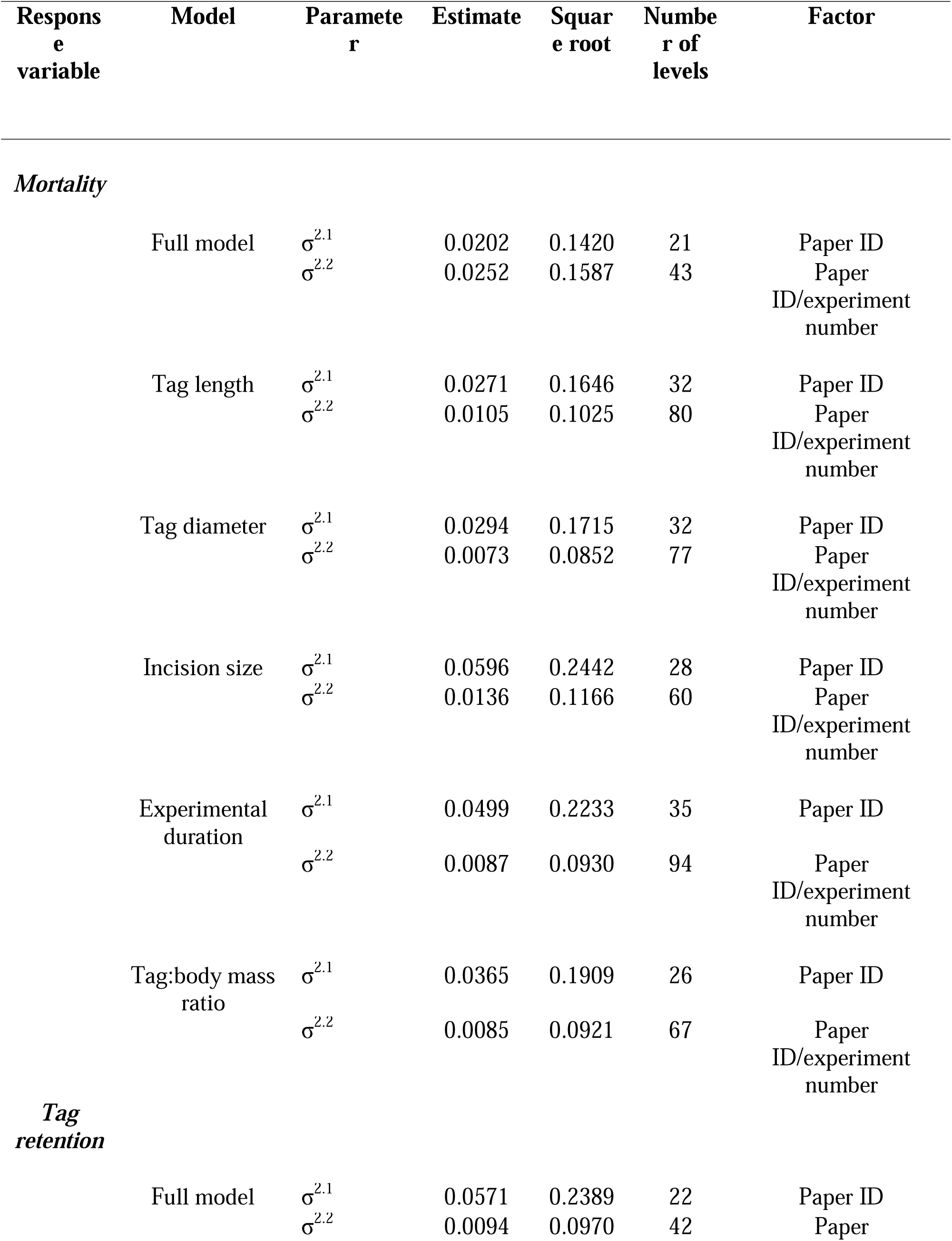
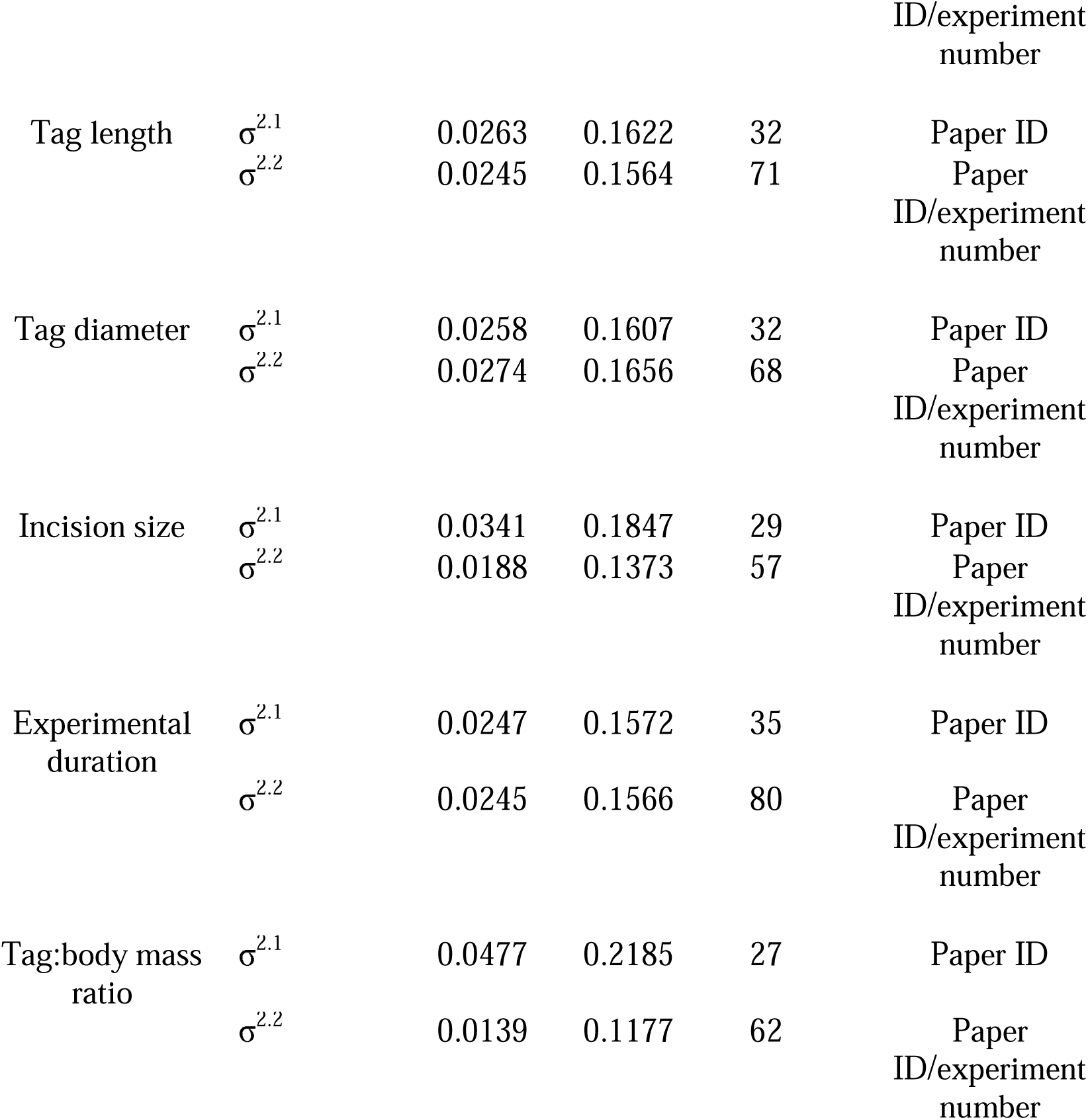
Meta-regression variance components for comparing tagging and experimental parameters against tag retention and mortality rates in fishes. Paper ID was treated as a random effect in the model while experimental number was nested within this random effect. Meta-regressions were conducted using proportional data transformed using a Freeman–Tukey double-arcsine transformation.

| Response variable | Model | Parameter | Estimate | Square root | Number of levels | Factor |
| --- | --- | --- | --- | --- | --- | --- |
| *Mortality* |  |  |  |  |  |  |
| | Full model | $\sigma^{2.1}$ | 0.0202 | 0.1420 | 21 | Paper ID |
| | | $\sigma^{2.2}$ | 0.0252 | 0.1587 | 43 | Paper ID/experiment number |
| | Tag length | $\sigma^{2.1}$ | 0.0271 | 0.1646 | 32 | Paper ID |
| | | $\sigma^{2.2}$ | 0.0105 | 0.1025 | 80 | Paper ID/experiment number |
| | Tag diameter | $\sigma^{2.1}$ | 0.0294 | 0.1715 | 32 | Paper ID |
| | | $\sigma^{2.2}$ | 0.0073 | 0.0852 | 77 | Paper ID/experiment number |
| | Incision size | $\sigma^{2.1}$ | 0.0596 | 0.2442 | 28 | Paper ID |
| | | $\sigma^{2.2}$ | 0.0136 | 0.1166 | 60 | Paper ID/experiment number |
| | Experimental duration | $\sigma^{2.1}$ | 0.0499 | 0.2233 | 35 | Paper ID |
| | | $\sigma^{2.2}$ | 0.0087 | 0.0930 | 94 | Paper ID/experiment number |
| | Tag:body mass ratio | $\sigma^{2.1}$ | 0.0365 | 0.1909 | 26 | Paper ID |
| | | $\sigma^{2.2}$ | 0.0085 | 0.0921 | 67 | Paper ID/experiment number |
| *Tag retention* |  |  |  |  |  |  |
| | Full model | $\sigma^{2.1}$ | 0.0571 | 0.2389 | 22 | Paper ID |
| | | $\sigma^{2.2}$ | 0.0094 | 0.0970 | 42 | Paper |

|  |  |  |  |  |  |
| --- | --- | --- | --- | --- | --- |
|  |  |  |  |  | ID/experiment<br>number |
| Tag length | $\sigma^{2.1}$ | 0.0263 | 0.1622 | 32 | Paper ID |
| | $\sigma^{2.2}$ | 0.0245 | 0.1564 | 71 | Paper<br>ID/experiment<br>number |
| Tag diameter | $\sigma^{2.1}$ | 0.0258 | 0.1607 | 32 | Paper ID |
| | $\sigma^{2.2}$ | 0.0274 | 0.1656 | 68 | Paper<br>ID/experiment<br>number |
| Incision size | $\sigma^{2.1}$ | 0.0341 | 0.1847 | 29 | Paper ID |
| | $\sigma^{2.2}$ | 0.0188 | 0.1373 | 57 | Paper<br>ID/experiment<br>number |
| Experimental<br>duration | $\sigma^{2.1}$ | 0.0247 | 0.1572 | 35 | Paper ID |
| | $\sigma^{2.2}$ | 0.0245 | 0.1566 | 80 | Paper<br>ID/experiment<br>number |
| Tag:body mass<br>ratio | $\sigma^{2.1}$ | 0.0477 | 0.2185 | 27 | Paper ID |
| | $\sigma^{2.2}$ | 0.0139 | 0.1177 | 62 | Paper<br>ID/experiment<br>number |

**Figure S1:**
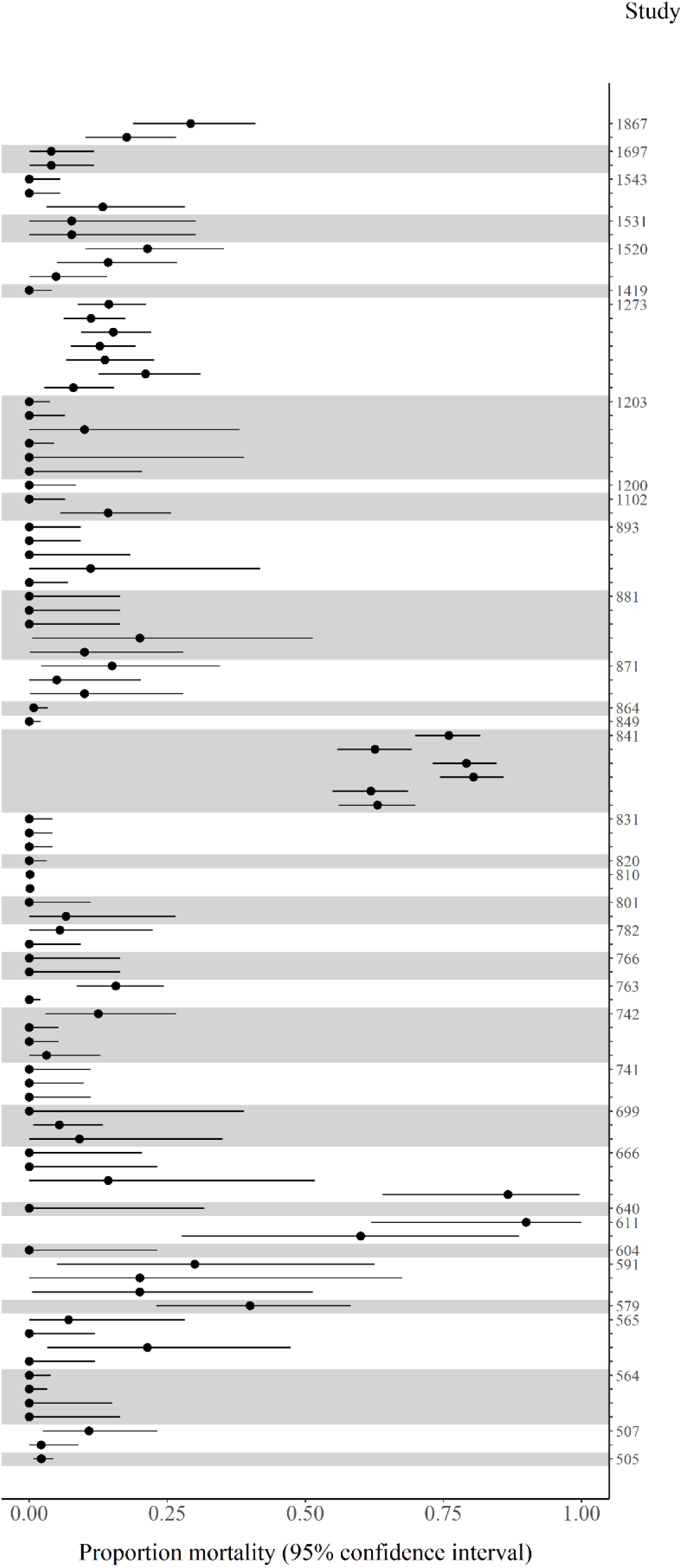
Forest plot depicting point estimates and the corresponding 95% confidence intervals for mortality proportions in tagged fish from journal articles collected in the metanalysis. Individual studies are represented by a coded number, which reference details can be found in Table S1. In some cases studies collected multiple estimates of tagging associated mortality which are represented by ticks following the study identifier.

**Figure S2:**
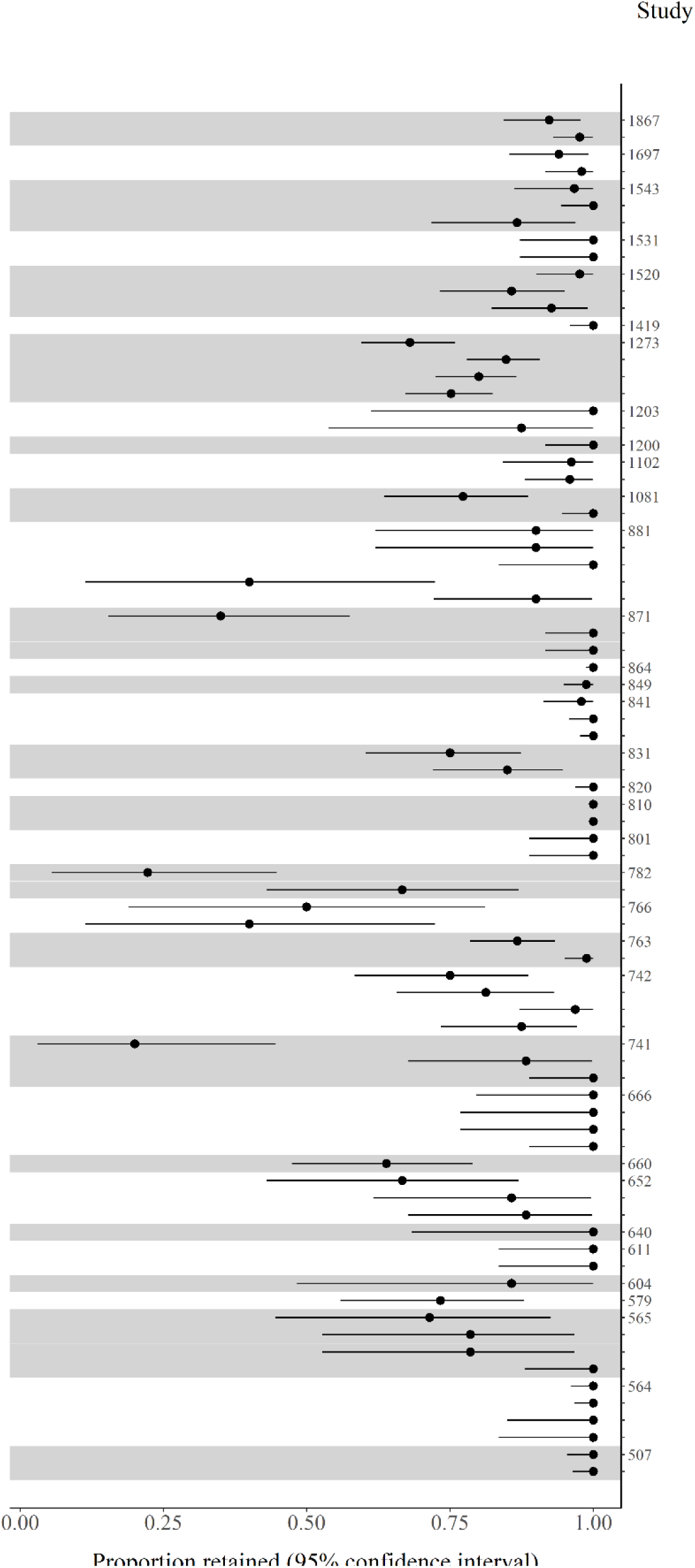
Forest plot depicting point estimates and the corresponding 95% confidence intervals for tag retention proportions in tagged fish from journal articles collected in the metanalysis. Individual studies are represented by a coded number, which reference details can be found in Table S1. In some cases studies collected multiple estimates of tagging associated mortality which are represented by ticks following the study identifier.

## Notes

https://github.com/mlaw27/Salmon-tag-retention-2023?search=1

### Competing Interest Statement

The authors have declared no competing interest.

## References

1. Crossin GT, Heupel MR, Holbrook CM, Hussey NE, Lowerre-Barbieri SK, Nguyen VM, et al. Acoustic telemetry and fisheries management. Ecol Appl. 2017;27:1031–49.

2. Heupel MR, Kessel ST, Matley JK, Simpfendorfer CA. Acoustic telemetry. CRC Press; 2018.

3. Donaldson MR, Hinch SG, Suski CD, Fisk AT, Heupel MR, Cooke SJ. Making connections in aquatic ecosystems with acoustic telemetry monitoring. Frontiers in Ecology and the Environment. 2014;12:565–73.

4. Raby GD, Vandergoot CS, Hayden TA, Faust MD, Kraus RT, Dettmers JM, et al. Does behavioural thermoregulation underlie seasonal movements in Lake Erie walleye? Canadian Journal of Fisheries and Aquatic Sciences. NRC Research Press; 2018;75:488–96.

5. Daly R, Filmalter JD, Daly CA, Bennett RH, Pereira MA, Mann BQ, et al. Acoustic telemetry reveals multi-seasonal spatiotemporal dynamics of a giant trevally Caranx ignobilis aggregation. Marine Ecology Progress Series. 2019;621:185–97.

6. Marsden JE, Blanchfield PJ, Brooks JL, Fernandes T, Fisk AT, Futia MH, et al. Using untapped telemetry data to explore the winter biology of freshwater fish. Rev Fish Biol Fisheries. 2021;31:115–34.

7. Childs A-R, Cowley PD, Næsje TF, Booth AJ, Potts WM, Thorstad EB, et al. Do environmental factors influence the movement of estuarine fish? A case study using acoustic telemetry. Estuarine, Coastal and Shelf Science. 2008;78:227–36.

8. Taylor MD, Fairfax AV, Suthers IM. The race for space: Using acoustic telemetry to understand density-dependent emigration and habitat selection in a released predatory fish. Reviews in Fisheries Science. Taylor & Francis; 2013;21:276–85.

9. Capra H, Plichard L, Bergé J, Pella H, Ovidio M, McNeil E, et al. Fish habitat selection in a large hydropeaking river: Strong individual and temporal variations revealed by telemetry. Science of the Total Environment. Elsevier; 2017;578:109–20.

10. Halfyard EA, Webber D, Del Papa J, Leadley T, Kessel ST, Colborne SF, et al. Evaluation of an acoustic telemetry transmitter designed to identify predation events. Börger L, editor. Methods Ecol Evol. 2017;8:1063–71.

11. Weinz AA, Matley JK, Klinard NV, Fisk AT, Colborne SF. Identification of predation events in wild fish using novel acoustic transmitters. Animal Biotelemetry. Springer; 2020;8:1–14.

12. Curtis JM, Johnson MW, Diamond SL, Stunz GW. Quantifying delayed mortality from barotrauma impairment in discarded red snapper using acoustic telemetry. Marine and Coastal Fisheries. Taylor & Francis; 2015;7:434–49.

13. Block BA, Whitlock R, Schallert RJ, Wilson S, Stokesbury MJW, Castleton M, et al. Estimating Natural Mortality of Atlantic Bluefin Tuna Using Acoustic Telemetry. Sci Rep. 2019;9:4918.

14. Flávio H, Kennedy R, Ensing D, Jepsen N, Aarestrup K. Marine mortality in the river? Atlantic salmon smolts under high predation pressure in the last kilometres of a river monitored for stock assessment. Fisheries Management and Ecology. Wiley Online Library; 2020;27:92– 101.

15. VillegasLRíos D, Freitas C, Moland E, Thorbjørnsen SH, Olsen EM. Inferring individual fate from aquatic acoustic telemetry data. Methods in Ecology and Evolution. Wiley Online Library; 2020;11:1186–98.

16. Welch DW, Melnychuk MC, Payne JC, Rechisky EL, Porter AD, Jackson GD, et al. In situ measurement of coastal ocean movements and survival of juvenile Pacific salmon. Proceedings of the National Academy of Sciences. National Acad Sciences; 2011;108:8708–13.

17. Wilson S, Hinch S, Drenner S, Martins E, Furey N, Patterson D, et al. Coastal marine and in-river migration behaviour of adult sockeye salmon en route to spawning grounds. Marine Ecology Progress Series. 2014;496:71–84.

18. Furey NB, Vincent SP, Hinch SG, Welch DW. Variability in migration routes influences early marine survival of juvenile salmon smolts. PLoS One. Public Library of Science San Francisco, CA USA; 2015;10:e0139269.

19. Freshwater C, Trudel M, Beacham TD, Godbout L, Neville C-EM, Tucker S, et al. Divergent migratory behaviours associated with body size and ocean entry phenology in juvenile sockeye salmon. Canadian Journal of Fisheries and Aquatic Sciences. NRC Research Press; 2016;73:1723–32.

20. Wagner GN, Cooke SJ, Brown RS, Deters KA. Surgical implantation techniques for electronic tags in fish. Reviews in Fish Biology and Fisheries. Springer; 2011;21:71–81.

21. Moore A, Russell I, Potter E. The effects of intraperitoneally implanted dummy acoustic transmitters on the behaviour and physiology of juvenile Atlantic salmon, Salmo salar L. Journal of fish biology. Wiley Online Library; 1990;37:713–21.

22. Bridger CJ, Booth RK. The effects of biotelemetry transmitter presence and attachment procedures on fish physiology and behavior. Reviews in Fisheries Science. Taylor & Francis; 2003;11:13–34.

23. Caputo M, O’Connor CM, Hasler CT, Hanson KC, Cooke SJ. Long-term effects of surgically implanted telemetry tags on the nutritional physiology and condition of wild freshwater fish. Diseases of Aquatic Organisms. 2009;84:35–41.

24. Ammann AJ, Michel CJ, MacFarlane RB. The effects of surgically implanted acoustic transmitters on laboratory growth, survival and tag retention in hatchery yearling Chinook salmon. Environmental Biology of Fishes. Springer; 2013;96:135–43.

25. Miller KM, Teffer A, Tucker S, Li S, Schulze AD, Trudel M, et al. Infectious disease, shifting climates, and opportunistic predators: cumulative factors potentially impacting wild salmon declines. Evolutionary Applications. Wiley Online Library; 2014;7:812–55.

26. Klinard NV, Halfyard EA, Fisk AT, Stewart TJ, Johnson TB. Effects of surgically implanted acoustic tags on body condition, growth, and survival in a small, laterally compressed forage fish. Transactions of the American Fisheries Society. Wiley Online Library; 2018;147:749–57.

27. Cooke SJ, Woodley CM, Brad Eppard M, Brown RS, Nielsen JL. Advancing the surgical implantation of electronic tags in fish: a gap analysis and research agenda based on a review of trends in intracoelomic tagging effects studies. Rev Fish Biol Fisheries. 2011;21:127–51.

28. Matley JK, Klinard NV, Barbosa Martins AP, Aarestrup K, Aspillaga E, Cooke SJ, et al. Global trends in aquatic animal tracking with acoustic telemetry. Trends in Ecology & Evolution. 2022;37:79–94.

29. Vollset KW, Lennox RJ, Thorstad EB, Auer S, Bär K, Larsen MH, et al. Systematic review and meta-analysis of PIT tagging effects on mortality and growth of juvenile salmonids. Rev Fish Biol Fisheries. 2020;30:553–68.

30. Wright DW, Stien LH, Dempster T, Oppedal F. Differential effects of internal tagging depending on depth treatment in Atlantic salmon: a cautionary tale for aquatic animal tag use. Current Zoology. Oxford University Press; 2019;65:665–73.

31. Collins AL, Hinch SG, Welch DW, Cooke SJ, Clark TD. Intracoelomic acoustic tagging of juvenile sockeye salmon: swimming performance, survival, and postsurgical wound healing in freshwater and during a transition to seawater. Transactions of the American Fisheries Society. Taylor & Francis; 2013;142:515–23.

32. Brunsdon EB, Daniels J, Hanke A, Carr J. Tag retention and survival of Atlantic salmon (Salmo salar) smolts surgically implanted with dummy acoustic transmitters during the transition from fresh to salt water. ICES Journal of Marine Science. Oxford University Press; 2019;76:2471–80.

33. Lacroix GL, Knox D, McCurdy P. Effects of implanted dummy acoustic transmitters on juvenile Atlantic salmon. Transactions of the American Fisheries Society. Taylor & Francis; 2004;133:211–20.

34. Honkanen HM, Rodger JR, Stephen A, Adams K, Freeman J, Adams CE. Counterintuitive migration patterns by Atlantic salmon Salmo salar smolts in a large lake. J Fish Biol. 2018;93:159–62.

35. Kennedy RJ, Rosell R, Millane M, Doherty D, Allen M. Migration and survival of Atlantic salmon Salmo salar smolts in a large natural lake. J Fish Biol. 2018;93:134–7.

36. R Core Team. R: A language and environment for statistical computing. [Internet]. Vienna, Austria: R Foudnation for Statistical Computing; 2021. Available from: https://www.R-project.org/

37. Therneau TM, Grambsch PM. Modeling Survival Data: Extending the Cox Model. 2000.

38. Therneau T. A Package for Survival Analysis in R [Internet]. 2021. Available from: <URL: https://CRAN.R-project.org/package=survival>.

39. Kassambara A, Kosinski M, Biecek P. survminer: Drawing Survival Curves using “ggplot2” [Internet]. 2021. Available from: https://CRAN.R-project.org/package=survminer

40. Crane DP, Killourhy CC, Clapsadl MD. Effects of three frozen storage methods on wet weight of fish. Fisheries Research. 2016;175:142–7.

41. Bates D, Mächler M, Bolker B, Walker S. Fitting Linear Mixed-Effects Models Using lme4. Journal of Statistical Software. 2015;67:1–48.

42. Lee S, Lee DK. What is the proper way to apply the multiple comparison test? Korean journal of anesthesiology. Korean Society of Anesthesiologists; 2018;71:353–60.

43. Lenth RV. emmeans: Estimated Marginal Means, aka Least-Squares Means [Internet]. 2023. Available from: https://CRAN.R-project.org/package=emmeans

44. Viechtbauer W. Conducting meta-analyses in R with the metafor package. Journal of statistical software. UCLA Statistics; 2010;36:1–48.

45. Freeman MF, Tukey JW. Transformations related to the angular and the square root. The Annals of Mathematical Statistics. JSTOR; 1950;607–11.

46. Viechtbauer W. Bias and efficiency of meta-analytic variance estimators in the random-effects model. Journal of Educational and Behavioral Statistics. Sage Publications Sage CA: Los Angeles, CA; 2005;30:261–93.

47. Benjamini Y, Hochberg Y. Controlling the false discovery rate: a practical and powerful approach to multiple testing. Journal of the Royal Statistical Society: series B (statistical methodology). Wiley Online Library; 1995;57:289–300.

48. Raue A, Kreutz C, Maiwald T, Bachmann J, Schilling M, Klingmüller U, et al. Structural and practical identifiability analysis of partially observed dynamical models by exploiting the profile likelihood. Bioinformatics. Oxford University Press; 2009;25:1923–9.

49. Welch D, Batten S, Ward B. Growth, survival, and tag retention of steelhead trout (O. mykiss) surgically implanted with dummy acoustic tags. Developments in Fish Telemetry. Springer; 2007. p. 289–99.

50. Chittenden CM, Butterworth KG, Cubitt KF, Jacobs MC, Ladouceur A, Welch DW, et al. Maximum tag to body size ratios for an endangered coho salmon (O. kisutch) stock based on physiology and performance. Environmental Biology of Fishes. Springer; 2009;84:129–40.

51. Rechisky EL, Welch DW. Surgical implantation of acoustic tags: Influence of tag loss and tag-induced mortality on free-ranging and hatchery-held spring Chinook (O. tschawytscha) smolts. PNAMP Special Publication: tagging, telemetry and marking neasures for monitoring fish populations—A compendium of new and recent science for use in informing technique and decision modalities: Pacific Northwest Aquatic Monitoring Partnership Special Publication. 2010;2:71–96.

52. Thorstad EB, Økland F, Westerberg H, Aarestrup K, Metcalfe JD. Evaluation of surgical implantation of electronic tags in European eel and effects of different suture materials. Marine and Freshwater Research. CSIRO Publishing; 2013;64:324–31.

53. Ward DL, Persons WR, Young KL, Stone DM, Vanhaverbeke DR, Knight WK. A laboratory evaluation of tagging-related mortality and tag loss in juvenile Humpback Chub. North American Journal of Fisheries Management. Taylor & Francis; 2015;35:135–40.

54. D’Arcy J, Kelly S, McDermott T, Hyland J, Jackson D, Bolton-Warberg M. Assessment of PIT tag retention, growth and post-tagging survival in juvenile lumpfish, Cyclopterus lumpus. Animal Biotelemetry. BioMed Central; 2020;8:1–9.

55. Jepsen N, Koed A, Thorstad EB, Baras E. Surgical implantation of telemetry transmitters in fish: how much have we learned? Aquatic telemetry. Springer; 2002. p. 239–48.

56. Sokolova IM. Energy-Limited Tolerance to Stress as a Conceptual Framework to Integrate the Effects of Multiple Stressors. Integrative and Comparative Biology. 2013;53:597–608.

57. Lawrence MJ, Godin J-GJ, Zolderdo AJ, Cooke SJ. Chronic Plasma Cortisol Elevation Does Not Promote Riskier Behavior in a Teleost Fish: A Test of the Behavioral Resiliency Hypothesis. Integrative Organismal Biology. 2019;1:obz009.

58. Romero LM, Dickens MJ, Cyr NE. The reactive scope model — A new model integrating homeostasis, allostasis, and stress. Hormones and Behavior. 2009;55:375–89.

59. Stobo WT, Fowler GM, Sinclair AF. Short-term tagging mortality of laboratory held juvenile Atlantic herring (Clupea h. harengus). Journal of Northwest Atlantic Fishery Science. Northwest Atlantic Fisheries Organization, NAFO; 1992;12.

60. Simard LG, Sotola VA, Marsden JE, Miehls S. Assessment of PIT tag retention and post-tagging survival in metamorphosing juvenile sea lamprey. Animal Biotelemetry. BioMed Central; 2017;5:1–7.

61. Bass AL, Stevenson CF, Porter AD, Rechisky EL, Furey NB, Healy SJ, et al. In situ experimental evaluation of tag burden and gill biopsy reveals survival impacts on migrating juvenile sockeye salmon. Can J Fish Aquat Sci. 2020;77:1865–9.

62. McKenna JE, Sethi SA, Scholten GM, Kraus J, Chalupnicki M. Acoustic tag retention and tagging mortality of juvenile cisco Coregonus artedi. Journal of Great Lakes Research. 2021;47:937–42.

63. Liss SA, Znotinas KR, Blackburn SE, Fischer ES, Hughes JS, Harnish RA, et al. From 95 to 59 millimetres: a new active acoustic tag size guideline for salmon. Can J Fish Aquat Sci. NRC Research Press; 2021;78:943–57.

64. Winter JD. Underwater biotelemetry. Fisheries techniques. American Fisheries Society; 1983;

65. Newton M, Barry J, Dodd JA, Lucas MC, Boylan P, Adams CE. Does size matter? A test of size-specific mortality in Atlantic salmon Salmo salar smolts tagged with acoustic transmitters. Journal of Fish Biology. 2016;89:1641–50.

66. Smircich MG, Kelly JT. Extending the 2% rule: the effects of heavy internal tags on stress physiology, swimming performance, and growth in brook trout. Animal Biotelemetry. 2014;2:16.

67. Brownscombe JW, Lédée EJI, Raby GD, Struthers DP, Gutowsky LFG, Nguyen VM, et al. Conducting and interpreting fish telemetry studies: considerations for researchers and resource managers. Rev Fish Biol Fisheries. 2019;29:369–400.

68. McArley T, Herbert N. Mortality, physiological stress and reflex impairment in sub-legal Pagrus auratus exposed to simulated angling. Journal of Experimental Marine Biology and Ecology. Elsevier; 2014;461:61–72.

69. Barton BA, Haukenes AH, Parsons BG, Reed JR. Plasma cortisol and chloride stress responses in juvenile walleyes during capture, transport, and stocking procedures. North American Journal of Aquaculture. Taylor & Francis; 2003;65:210–9.

70. Danylchuk AJ, Suski CD, Mandelman JW, Murchie KJ, Haak CR, Brooks AM, et al. Hooking injury, physiological status and short-term mortality of juvenile lemon sharks (Negaprion bevirostris) following catch-and-release recreational angling. Conservation Physiology. Oxford University Press; 2014;2:cot036.

71. Mandelman JW, Skomal GB. Differential sensitivity to capture stress assessed by blood acid–base status in five carcharhinid sharks. J Comp Physiol B. 2009;179:267–77.

72. Lawrence MJ, Jain-Schlaepfer S, Zolderdo AJ, Algera DA, Gilmour KM, Gallagher AJ, et al. Are 3 minutes good enough for obtaining baseline physiological samples from teleost fish? Can J Zool. 2018;96:774–86.

73. Pickering AD, Pottinger TG, Christie P. Recovery of the brown trout, Salmo trutta L., from acute handling stress: a time-course study. J Fish Biology. 1982;20:229–44.

74. Foo J, Lam T. Serum cortisol response to handling stress and the effect of cortisol implantation on testosterone level in the tilapia, Oreochromis mossambicus. Aquaculture. Elsevier; 1993;115:145–58.

75. Martinelli TL, Hansel H, Shively R. Growth and physiological responses to surgical and gastric radio transmitter implantation techniques in subyearling chinook salmon (Oncorhynchus tshawytscha). Springer; 1998. p. 79–87.

76. Baker DW, Peake SJ, Kieffer JD. The effect of capture, handling, and tagging on hematological variables in wild adult lake sturgeon. North American Journal of Fisheries Management. Taylor & Francis; 2008;28:296–300.

77. Davis KB, Griffin BR. Physiological responses of hybrid striped bass under sedation by several anesthetics. Aquaculture. Elsevier; 2004;233:531–48.

78. Yousaf MN, Røn Ø, Hagen PP, McGurk C. Monitoring fish welfare using heart rate bio-loggers in farmed Atlantic salmon (Salmo salar L.): An insight into the surgical recovery. Aquaculture. Elsevier; 2022;555:738211.

79. Lower N, Moore A, Scott AP, Ellis T, James JD, Russell IC. A non-invasive method to assess the impact of electronic tag insertion on stress levels in fishes. Journal of Fish Biology. 2005;67:1202–12.

80. Jepsen N, Davis L, Schreck C, Siddens B. The physiological response of chinook salmon smolts to two methods of radio-tagging. Transactions of the American Fisheries Society. Taylor & Francis; 2001;130:495–500.

81. Lawrence MJ, Raby GD, Teffer AK, Jeffries KM, Danylchuk AJ, Eliason EJ, et al. Best practices for non-lethal blood sampling of fish via the caudal vasculature. Journal of Fish Biology. 2020;97:4–15.

82. Holland KN. A perspective on billfish biological research and recommendations for the future. Marine and Freshwater Research. CSIRO Publishing; 2003;54:343–7.

83. Chateau O, Wantiez L. Post-release activity of three coral reef fish species in a marine reserve: analysis and recommendations for telemetry studies. Environmental Biology of Fishes. Springer; 2021;104:15–26.

84. Jepsen N, Schreck C, Clements S, Thorstad EB. A brief discussion on the 2% tag/bodymass rule of thumb. Aquatic telemetry: advances and applications. 2005;255–9.

85. Økland F, Thorstad EB. Recommendations on size and position of surgically and gastrically implanted electronic tags in European silver eel. Animal Biotelemetry. Springer; 2013;1:1–6.

86. Leroy B, Scutt Phillips J, Potts J, Brill RW, Evans K, Forget F, et al. Recommendations towards the establishment of best practice standards for handling and intracoelomic implantation of data-storage and telemetry tags in tropical tunas. Animal Biotelemetry. BioMed Central; 2023;11:1–17.

87. Knights BC, Lasee BA. Effects of implanted transmitters on adult bluegills at two temperatures. Transactions of the American Fisheries Society. Wiley Online Library; 1996;125:440–9.

88. Holbrook SC, Byars WD, Lamprecht SD, Leitner JK. Retention and physiological effects of surgically implanted telemetry transmitters in Blue Catfish. North American Journal of Fisheries Management. Taylor & Francis; 2012;32:276–81.

89. Meyer CG, Honebrink RR. Transintestinal expulsion of surgically implanted dummy transmitters by bluefin trevally—implications for long-term movement studies. Transactions of the American Fisheries Society. Taylor & Francis; 2005;134:602–6.

90. Marty GD, Summerfelt RC. Pathways and mechanisms for expulsion of surgically implanted dummy transmitters from channel catfish. Transactions of the American Fisheries Society. Taylor & Francis; 1986;115:577–89.

91. Anderson JM, Rodriguez A, Chang DT. Foreign body reaction to biomaterials. Elsevier; 2008. p. 86–100.

92. Carnicer-Lombarte A, Chen S-T, Malliaras GG, Barone DG. Foreign body reaction to implanted biomaterials and its impact in nerve neuroprosthetics. Frontiers in Bioengineering and Biotechnology. Frontiers; 2021;271.

93. Ward WK. A review of the foreign-body response to subcutaneously-implanted devices: the role of macrophages and cytokines in biofouling and fibrosis. Journal of diabetes science and technology. SAGE Publications; 2008;2:768–77.

94. Onuki Y, Bhardwaj U, Papadimitrakopoulos F, Burgess DJ. A review of the biocompatibility of implantable devices: current challenges to overcome foreign body response. Journal of diabetes science and technology. SAGE Publications; 2008;2:1003–15.

95. Bostan H, Karakaya M, Demir M, Çağdir A, Hanci V. A case of surgical instrument left in the abdomen and taken out of the transverse colon. Hippokratia. Hippokratio General Hospital of Thessaloniki; 2014;18:77.

96. Nuovo J, Sweha A. Keloid formation from levonorgestrel implant (Norplant System) insertion. The Journal of the American Board of Family Practice. Am Board Family Med; 1994;7:152–4.

97. Lucas M. Effects of implanted dummy transmitters on mortality, growth and tissue reaction in rainbow trout, Salmo gairdneri Richardson. Journal of Fish biology. Wiley Online Library; 1989;35:577–87.

98. Baras E, Westerloppe L. Transintestinal expulsion of surgically implanted tags by African catfish Heterobranchus longifilis of variable size and age. Transactions of the American Fisheries Society. Taylor & Francis; 1999;128:737–46.

99. Gheorghiu C, Hanna J, Smith JW, Smith DS, Wilkie MP. Encapsulation and migration of PIT tags implanted in brown trout (Salmo trutta L.). Aquaculture. Elsevier; 2010;298:350–3.

100. Gries G, Letcher B. Tag retention and survival of age-0 Atlantic salmon following surgical implantation with passive integrated transponder tags. North American Journal of Fisheries Management. Taylor & Francis; 2002;22:219–22.

101. Foldvik A, Kvingedal E. Long-term PIT tag retention rates in Atlantic salmon (Salmo salar). Animal Biotelemetry. Springer; 2018;6:1–4.

102. Robinson RR, Notch J, McHuron A, Logston R, Pham T, Ammann AJ. The effects of water temperature, acoustic tag type, size at tagging, and surgeon experience on juvenile Chinook salmon (Oncorhynchus tshawytscha) tag retention and growth. Animal Biotelemetry. 2021;9:22.

103. Fischer ES, Blackburn SE, Liss SA, Hughes JS, Li H, Deng ZD. How small can we go? Evaluating survival, tag retention, and growth of juvenile Chinook salmon implanted with a new acoustic microtag. North American Journal of Fisheries Management. Wiley Online Library; 2019;39:1329–36.

104. Huusko R, Huusko A, MäkiLPetäys A, Orell P, Erkinaro J. Effects of tagging on migration behaviour, survival and growth of hatcheryLreared Atlantic salmon smolts. Fisheries Management and Ecology. Wiley Online Library; 2016;23:367–75.

105. D’Amico TW, Winkelman DL, Swarr TR, Myrick CA. Retention of passive integrated transponder tags in a smallLbodied catfish. North American Journal of Fisheries Management. Wiley Online Library; 2021;41:187–95.

106. Deters KA, Brown RS, Carter KM, Boyd JW, Eppard MB, Seaburg AG. Performance assessment of suture type, water temperature, and surgeon skill in juvenile Chinook salmon surgically implanted with acoustic transmitters. Transactions of the American Fisheries Society. Taylor & Francis; 2010;139:888–99.

107. Jonsson B, Forseth T, Jensen A, Næsje T. Thermal performance of juvenile Atlantic Salmon, Salmo salar L. Functional Ecology. Wiley Online Library; 2001;15:701–11.

108. Anderson JM, Jiang S. Implications of the acute and chronic inflammatory response and the foreign body reaction to the immune response of implanted biomaterials. The Immune Response to Implanted Materials and Devices: The Impact of the Immune System on the Success of an Implant. Springer; 2017;15–36.

109. Sloman KA, Bouyoucos IA, Brooks EJ, Sneddon LU. Ethical considerations in fish research. J Fish Biol. 2019;94:556–77.

110. Greenstreet SPR, Morgan RIG. The effect of ultrasonic tags on the growth rates of Atlantic salmon, Salmo salar L., parr of varying size just prior to smolting. Journal of Fish Biology. 1989;35:301–9.

111. Brown RS, Geist DR, Deters KA, Grassell A. Effects of surgically implanted acoustic transmitters> 2% of body mass on the swimming performance, survival and growth of juvenile sockeye and Chinook salmon. Journal of Fish Biology. Wiley Online Library; 2006;69:1626–38.

112. Brown RS, Harnish RA, Carter KM, Boyd JW, Deters KA, Eppard MB. An evaluation of the maximum tag burden for implantation of acoustic transmitters in juvenile Chinook salmon. North American Journal of Fisheries Management. Taylor & Francis; 2010;30:499–505.

## References

113. Baras E, Westerloppe L. Transintestinal expulsion of surgically implanted tags by African catfish Heterobranchus longifilis of variable size and age. Trans Am Fish Soc. 1999;128:737– 46.

114. Brundson EB, Daniels J, Hanke A, Carr J. Tag retention and survival of Atlantic salmon (Salmo salar) smolts surgically implanted with dummy acoustic transmitters during the transition from fresh to salt water. ICES J mar Sci. 2019;76:2471–80.

115. Butler GL, Mackay B, Rowland SJ, Pease BC. Retention of intra-peritoneal transmitters and post-operative recovery of four Australian native fish species. Mar Freshwater Res. 2009;60:361.

116. Childs AR, Næsje TF, Cowley PD. Long-term effects of different-sized surgically implanted acoustic transmitters on the sciaenid Arygyrosomus japonicus: breaking the 2% tag-to-body mass rule. Mar Freshwater Res. 2011;62:432.

117. Cook KV, Brown RS, Daniel Deng Z, Klett RS, Li H, Seaburg AG, et al. A comparison of implantation methods for large PIT tags or injectable acoustic transmitters in juvenile Chinook salmon. Fish Res. 2014;154:213–23.

118. Daniel AJ, Hicks BJ, Ling N, David BO. Acoustic and radio-transmitter retention in common carp (Cyprinus carpio) in New Zealand. Mar Freshwater Res. 2009;60:328.

119. Ellis T, Buckel J, Hightower J. Winter severity influences spotted seatrout mortality in a southeast US estuarine system. Mar Ecol Prog Ser. 2017;564:145–61.

120. Fabrizio MC, Pessutti JP. Long-term effects and recovery from surgical implantation of dummy transmitters in two marine fishes. Journal of Experimental Marine Biology and Ecology. 2007;351:243–54.

121. Frost DA, McComas RL, Sandford BP. The effects of a surgically implanted microacoustic tag on growth and survival in subyearling fall chinook salmon. Trans Am Fish Soc. 2010;139:1192–7.

122. Gerber KM, Mather ME, Smith JM, Peterson ZJ. Evaluation of a field protocol for internally-tagging fish predators using difficult-to-tag ictalurid catfish as examples. Fish Res. 2019;209:58–66.

123. Halfyard EA, Webber D, Del Papa J, Leadley T, Kessel ST, Colborne SF, et al. Evaluation of an acoustic telemetry transmitter designed to identify predation events. Börger L, editor. Methods Ecol Evol. 2017;8:1063–71.

124. Harnish R, Deters K, Ham K, Deng Z, Li H, Rayamajhi B, et al. Survival of wild Hanford Reach and Priest Rapids hatchery fall chinook salmon juveniles in the Columbia River: Predation implications. Pacific Northwest National Labratory; 2014 p. 68. Report No.: PNNL-SA-23719.

125. Jepsen N, Larsen MH, Aarestrup K. Performance of fast absorbable sutures and histo-glue for closing incisions in brown trout. Trans Am Fish Soc. 2017;146:1233–7.

126. Karam AP, Kesner BR, Marsh PC. Acoustic telemetry to assess post-stocking dispersal and mortality of razorback sucker Xyrauchen texanus. J Fish Biol. 2008;73:719–27.

127. Kennedy RJ, Evans D, Allen M. LongLJterm retention of dummy acoustic transmitters in adult brown trout. J Fish Biol. 2020;97:1281–4.

128. Koeck B, Gudefin A, Romans P, Loubet J, Lenfant P. Effects of intracoelomic tagging procedure on white seabream (Diplodus sargus) behavior and survival. J Exp Mar Biol Ecol. 2013;440:1–7.

129. Loher T, Rensmeyer R. Physiological responses of Pacific halibut, Hippoglossus stenolepis, to intracoelomic implantation of electronic archival tags, with a review of tag implantation techniques employed in flatfishes. Rev Fish Biol Fish. 2011;21:97–115.

130. Luo H, Duan X, Wang S, Liu S, Chen D. Effects of surgically implanted dummy ultrasonic transmitters on growth, survival and transmitter retention of bighead carp Hypophthalmichthys nobilis. Environ Biol Fish. 2015;98:1131–9.

131. McKenna JE, Sethi SA, Scholten GM, Kraus J, Chalupnicki M. Acoustic tag retention and tagging mortality of juvenile cisco Coregonus artedi. J. Great Lakes Res. 2021;47:937– 42.

132. Melnychuk MC, Welch DW, Walters CJ, Christensen V. Riverine and early ocean migration and mortality patterns of juvenile steelhead trout (Oncorhynchus mykiss) from the Cheakamus River, British Columbia. Hydrobiologia. 2007;582:55–65.

133. Mueller R, Liss S, Deng ZD. Implantation of a new micro acoustic tag in juvenile Pacific lamprey and American eel. JoVE. 2019;59274.

134. Murchie KJ, Danylchuk AJ, Cooke SJ, O’Toole AC, Shultz A, Haak C, et al. Considerations for tagging and tracking fish in tropical coastal habitats: lessons from bonefish, barracuda, and sharks tagged with acoustic transmitters. In: Adams NS, Beeman JW, Eiler JH, editors. Telemetry techniques: A user guide for fisheries research. American Fisheries Society; 2012. p. 389–412.

135. Neves V, Silva D, Martinho F, Antunes C, Ramos S, Freitas V. Assessing the effects of internal and external acoustic tagging methods on European flounder Platichthys flesus. Fish Res. 2018;206:202–8.

136. Neville CM, Beamish RJ, Chittenden CM. Poor survival of acoustically-tagged juvenile chinook salmon in the Strait of Georgia, British Columbia, Canada. Trans Am Fish Soc. 2015;144:25–33.

137. Robinson RR, Notch J, McHuron A, Logston R, Pham T, Ammann AJ. The effects of water temperature, acoustic tag type, size at tagging, and surgeon experience on juvenile

138. Chinook salmon (Oncorhynchus tshawytscha) tag retention and growth. Anim Biotelem. 2021;9:22.

139. Rożyński M, Kapusta A, Demska-Zakęś K, Ziomek E, Szczerbowski A, Stawecki K, et al. Impact of two telemetry transmitter implantation incision suturing methods on the physiological state and condition of perch (Perca fluviatilis). Arch Pol Fish. 2017;25:89–101.

140. Sandstrom PT, Ammann AJ, Michel C, Singer G, Chapman ED, Lindley S, et al. Growth, survival, and tag retention of steelhead trout (Oncorhynchus mykiss) and its application to survival estimates. Environ Biol Fish. 2013;96:145–64.

141. Scheffel TK, Hightower JE, Buckel JA, Krause JR, Scharf FS. Coupling acoustic tracking with conventional tag returns to estimate mortality for a coastal flatfish with high rates of emigration. Can J Fish Aquat Sci. 2020;77:1–22.

142. Seitz AC, Norcross BL, Payne JC, Kagley AN, Meloy B, Gregg JL, et al. Feasibility of surgically implanting acoustic tags into Pacific herring. Trans Am Fish Soc. 2010;1391288–91.

143. Smircich MG, Kelly JT. Extending the 2% rule: the effects of heavy internal tags on stress physiology, swimming performance, and growth in brook trout. Anim Biotelem. 2014;2:16.

144. Stakėnas S, Copp GH, Scott DM. Tagging effects on three non-native fish species in England (Lepomis gibbosus, Pseudorasbora parva, Sander lucioperca) and of native Salmo trutta. Ecol Freshw Fish. 2009;18:167–76.

145. Thorstad EB, Kerwath SE, Attwood CG, Økland F, Wilke CG, Cowley PD, et al. Long-term effects of two sizes of surgically implanted acoustic transmitters on a predatory marine fish (Pomatomus saltatrix). Mar Freshwater Res. 2009;60:183.

146. Tsitrin E, McLean MF, Gibson AJF, Hardie DC, Stokesbury MJW. Feasibility of using surgical implantation methods for acoustically tagging alewife (Alosa pseudoharengus) with V5 acoustic transmitters. PLoS ONE. 2020;15:e0241118.

147. Urbaniak TJ, Barnes ME, Davis JL. Acoustic transmitters impact rainbow trout growth in a competitive environment. Open Fish Sci J. 2016;9:37–44.

148. Weiland MA, Ploskey GR, Hughes JS, Woodley CM, Deng Z, Carlson TJ, et al. Monitoring of juvenile yearling chinook salmon and steelhead survival and passage at John Day Dam, Spring 2010. Pacific Northwest National Laboratory; 2012 Nov p. PNNL-22015, 1097980. Report No.: PNNL-22015, 1097980.

149. Welch DW, Batten SD, Ward BR. Growth, survival, and tag retention of steelhead trout (O. mykiss) surgically implanted with dummy acoustic tags. Hydrobiologia. 2007;582:289– 99.

150. Woodley CM, Carpenter SM, Carter KM, Wagner KA, Royer IM, Knox KM, et al. Surgically implanted JSATS micro-acoustic transmitters effects on juvenile chinook salmon and steelhead tag expulsion and survival, 2010. Pacific Northwest National Laboratory; 2011 Sep p. PNNL-20857, 1028581. Report No.: PNNL-20857, 1028581.

151. Yasuda T, Nagano N, Kitano H, Ohga H, Sakai T, Ohshimo S, et al. Tag attachment success can be temperature dependent: a case study of the chub mackerel Scomber japonicus . Anim Biotelem. 2015;3:48.

152. Zemeckis DR, Hoffman WS, Dean MJ, Armstrong MP, Cadrin SX. Spawning site fidelity by Atlantic cod (Gadus morhua) in the Gulf of Maine: implications for population structure and rebuilding. ICES J Mar Sci. 2014;71:1356–65.

